# IL-32 drives inflammatory responses in IFN-γ primed human macrophages via a Myddosome-dependent pathway and is elevated in COVID-19

**DOI:** 10.1101/2025.05.29.656795

**Authors:** Ana Ramón-Vázquez, Agnieszka Skowyra, Tadhg Crowley, Janaki Velmurugan, Ciaran Lee, Andrew J. Lindsay, Jerzy A. Woznicki, Panagiota Stamou, Ornella Nelson, Ian B. Jeffery, Amanda J. Lohan, Werner C. Albrich, Liam O’Mahony, Silvia Melgar, Fergus Shanahan, Xiaohong Cao, Michael Macoritto, Ramkrishna Sadhukhan, Marc C. Levesque, Bradford L. McRae, Melissa Matzelle, Ken Nally

## Abstract

IFN-γ is secreted by multiple lymphoid subsets in response to antigen stimulation and can reprogram and prime macrophages epigenetically and transcriptionally to increase responses to inflammatory stimuli such as LPS, IL-1β or TNF-α. IFN-γ-driven M1-like inflammatory macrophage states are shared across human immune-mediated inflammatory diseases (IMIDs) and while IFN-γ is nonredundant for defense to intracellular pathogens it is unclear if this is also the case in IMIDs. To identify additional secreted ligands which could prime and induce M1-like macrophages we screened >600 human proteins in human primary macrophages. Using complementary functional genomics approaches, we discovered that IL-32β induced an M1-like inflammatory state in non-primed and IFN-γ-primed macrophages. IL-32β induced signaling, transcriptional, tolerance, cross-tolerance, and inflammatory responses in macrophages which were MyD88, IRAK1 and Myddosome-dependent. These responses to host IL-32β were similar to yet distinct from, those induced by microbial LPS. IL-32 protein was elevated in serum from patients with severe COVID-19 and IL-32β together with IFN-γ were expressed by T cells and induced a macrophage transcriptional response which was shared by monocytes and macrophages in mild and severe COVID-19.

## Introduction

IFN-γ is a cytokine produced and secreted by multiple lymphoid subsets (CD4^+^T_H_1, CD4^+^T_H_1*, CD8^+^, NK, NKT and ILC1 cells) in response to antigen stimulation by antigen presenting cells. While potentially all human cells can respond to IFN-γ; monocytes, monocyte-derived macrophages and tissue resident macrophages are the major targets of lymphoid cells that produce IFN-γ^1^. Once secreted, IFN-γ binds to the IFNGR1/IFNGR2 receptor complex on target cells leading to the activation of tyrosine kinases, Janus kinase 1 and 2 (JAK1/JAK2), which in turn facilitate recruitment, tyrosine 701 (Y701) phosphorylation and activation of signal transducer and activator of transcription 1 (STAT1)^2^. IFN-γ transcriptionally reprograms macrophages via the JAK1/2-STAT1 signaling pathway leading to the induction of hundreds of interferon stimulated genes (ISGs) which code for antimicrobial, cytotoxic, immunomodulatory and inflammatory effector proteins. IFN-γ also primes macrophages to respond more robustly and synergistically to secondary stimulation with other immune and inflammatory stimuli such as LPS, IL-1β or TNF-α^1,3-5^. These effects of IFN-γ on macrophages are underpinned by JAK1/2-STAT1 driven transcriptional, epigenetic and metabolic reprogramming leading to a transcriptional memory or trained immunity state which can persist for months^4^. The key molecular mechanisms underpinning the phenomenon of priming include modification of the epigenomic landscape through chromatin remodeling and deposition of histone modifications that increase or decrease chromatin accessibility for transcription factors in a gene-specific manner. In addition, upon stimulation of cells with IFN-γ, STAT1 relocates to non-canonical binding sites in promoters and enhancers of non-classical interferon-stimulated genes (ISGs) such as the signature cytokines TNF-α, IL-6, IL-12/23p40 and IL-10 which are used to evaluate inflammatory, priming, innate immune training, and tolerance responses in macrophages. There is an enrichment of IRF and NF-κB DNA binding motifs in these regions of non-canonical STAT1 binding, which can be explained by the interaction of STAT1 with IRF-containing protein complexes^4,6,7^.

The term human immune[mediated inflammatory diseases (IMID) define a group of incurable diseases of uncertain etiology that share common features, mainly relapsing and remitting chronic inflammation and immune dysregulation, but that also present unique characteristics, for example clinical phenotypes or therapeutic response profiles^8–10^. Examples include the inflammatory bowel diseases (IBD), Crohn’s disease (CD) and ulcerative colitis (UC), systemic lupus erythematosus (SLE), rheumatoid arthritis (RA), psoriasis, psoriatic arthritis, ankylosing spondylitis, asthma and neurological diseases such as multiple sclerosis (MS). Profiling of cell types and cell states across IMID tissues using single cell RNA sequencing (scRNA-seq) has demonstrated both shared and disease selective features in terms of changes in cell composition, immunopathogenic cell states, cell-cell interactions and responses to treatment, highlighting that the characterization, target identification and therapeutic targeting of IMIDs should be approached in a tissue and disease agnostic way^11^. During the last 10 years, scRNA-seq analysis has led to the identification and characterization of expanded inflammatory fibroblast, monocyte and macrophage cell states shared across multiple IMIDs^11,12^. Recently it has become clear that an IFN inflammatory signature in macrophages (and other cell types) is a common shared transcriptional phenotype and cell state associated with multiple IMIDs, infectious diseases such COVID-19 and inadequate therapeutic responses to anti-TNFs in IBD^5,13–19^. Interestingly, in the context of RA, CD, UC and severe COVID-19, TNF-α was identified as a cytokine that could co-operate with IFN-γ to drive a CXCL10^+^CCL2^+^ macrophage cell state which was like the transcriptional phenotype of macrophages expanded in these diseases^5^.

While IFN-γ is an essential and nonredundant cytokine for human host defense to intracellular pathogens it is unclear if this is also the case for its role in priming M1-like macrophages and driving inflammation and immunopathogenesis in IMIDs^1,7^. In this study, we investigated if other human secreted proteins were redundant with IFN-γ in terms of its priming effects on human macrophages and/or could co-operate and synergize with IFN-γ to drive M1-like inflammatory macrophage cell states. We found that IL-32β and IL-32γ but not IL-32α are secreted proteins which can co-operate with IFN-γ to drive macrophage inflammatory responses and that IL-32β does this in macrophages via a Myddosome signaling pathway. Overall, our results support an immunopathogenic model whereby T cell secreted IFN-γ and IL-32β mediate cell-cell communication between T cells and macrophages and co-operate to drive inflammatory macrophage responses in severe COVID-19.

## Results

### Human macrophage immunoinflammatory screens identify secreted ligands with IFN-γ-like priming and M1-like inflammatory activity

Given the apparent essential and nonredundant role of IFN-γ for human host defense to intracellular pathogens and its contribution to inflammation and immunopathogenesis in IMIDs, it is important to discover if other ligands have IFN-γ-like activity or can co-operate with IFN-γ to augment inflammatory macrophage responses^1^. To identify IMID relevant ligands, we compiled a database of 1,516 Crohn’s disease (CD) relevant genes from five publicly available gene expression datasets. These 1,516 genes code for 864 secreted ligands, 652 receptors and together accounted for 2,422 literature supported ligand-receptor pairs expressed in CD gut mucosal tissue (see methods section for details). Of these 1,516 genes, 601 recombinant proteins were sourced commercially to generate a protein library of secreted ligands. This library was screened in macrophage-based phenotypic screens to identify ligands with M1-like inflammatory activity similar to that of IFN-γ and the TLR4 ligand LPS, or anti-inflammatory activity like that of IL-10. Screens consisted of four different experimental workflows and production of signature cytokines (TNF-α, IL-6, IL-12/23p40 and IL-10) - used to evaluate inflammatory, priming, innate immune training, and tolerance responses in macrophages - were used as readouts (Figures 1A and S1A). Workflows were designed to identify ligands with effects on macrophages similar to that of (i) LPS (LPS-like or pro-inflammatory activity), (ii) IFN-γ (IFN-γ-like priming effect to LPS or IFN-γ-like activity), (iii) IFN-γ + LPS (LPS-like effect in IFN-γ-primed cells or M1-like activity), or (iv) IL-10 (IL-10-like effect or anti-inflammatory activity). Primary screens identified potential candidates after applying analysis filters (see methods section) (Figures 1A, S1A and Table S1). This initial list of ligands was revised to include ligands with Z scores below the set threshold but with known relevance to macrophage biology generating a list of 82 proteins for secondary screens using the same experimental workflows (Figures 1A, S1B and Table S2). Ligands that induced at least one of five cytokine readouts from TNF-α, IL-6, IL-12/23p40, IL-27 and IL-10 in secondary screens are shown (Figure S1B). The following protein ligands were identified and shortlisted from these screens for validation; IL-32γ and HSP90AA1 in the pro-inflammatory and M1-like screen, CSF2 and IL-3 in the IFN-γ-like screen and type-I IFNs in the IL-10-like screen.

**Figure 1.**
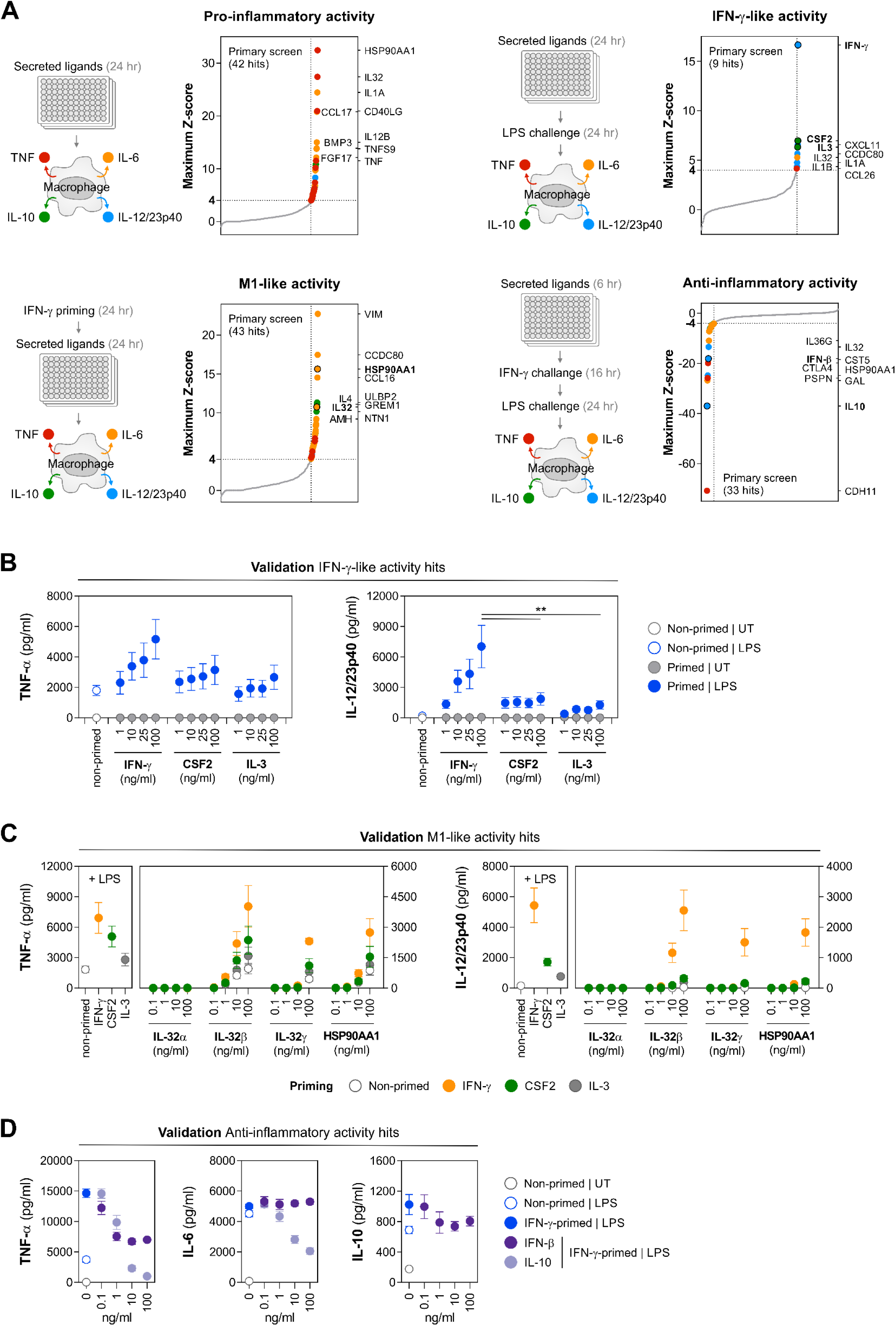
Immunoinflammatory screens in human macrophages identify secreted ligands with IFN-γ-like (CSF2, IL-3) and M1-like (IL-32β/γ, HSP90AA1) inflammatory activities. Also see Figure S1. (A) Schematics of experimental workflows and Z-score distribution graphs for hits from four different immunoinflammatory phenotypic screens performed in primary human macrophages with a library of 601 recombinant secreted ligands (see main text for details). Selected hits for further validation are highlighted in bold. (B) Validation of macrophage cytokine responses to priming effects of IFN-γ-like ligands. (C) Validation of M1-like ligands in non-primed, IFN-γ-primed, CSF2-primed and IL-3 primed macrophages. (D) Validation of macrophage cytokine responses to the anti-inflammatory ligand IFN-β. Statistical analysis: (B) represented data are mean values ±SEM of n = 3 independent experiments performed on monocyte-derived macrophages from one donor, one-way ANOVA with Tukey’s multiple comparisons test (** p < 0.005). LPS was used as positive control at 50 ng/mL in all cases.

To validate the IFN-γ-like ligands CSF2 (GM-CSF) and IL-3, we compared their ability to prime macrophages to LPS with the priming effects of IFN-γ in concentration-response experiments. Like IFN-γ, priming of macrophages with CSF2 and IL-3 could augment production of IL-6 and TNF-α by LPS relative to effects of LPS stimulation alone (Figure S1C and 1B). However, the priming effects of CSF2 and IL-3 were different to IFN-γ in terms of augmenting LPS-induced production of the cytokine subunit IL-12/23p40 and suppressing LPS-induced production of IL-10 relative to effects of LPS stimulation alone (Figure 1B and S1C). While CSF2 and IL-3 could prime macrophages for LPS-induced IL-12/23p40 production, the magnitude of the priming effect was much less and there was no concentration dependent priming response when compared to IFN-γ (Figure 1B). Similar to previous reports^20^, IFN-γ priming could reduce LPS-induced IL-10 production compared to LPS stimulation alone, however, CSF2 and IL-3 priming lacked this suppressive effect on LPS-induced IL-10 production (Figure S1C).

The IL-32 gene has 35 transcript variants (ensemble.org) of which 30 potentially encode proteins. Of these, nine protein isoforms have been validated experimentally and four isoforms IL-32α, IL-32β, IL-32γ and IL-32ε have been characterized most extensively in the literature. The IL-32γ isoform used in our macrophage screens was reported to be the most active isoform^21–23^. Therefore, we validated the potential pro-inflammatory activity of IL-32α, IL-32β, IL-32γ and another hit HSP90AA1, in non-primed macrophages and macrophages primed with IFN-γ, CSF2 and IL-3 (Figures 1C and S1D). Of the three IL-32 isoforms tested, IL-32β induced the most robust cytokine responses in non-primed and primed macrophages in a concentration-dependent manner. It was also more active at 10 ng/mL than IL-32γ with the greatest magnitude of response seen for TNF-α, IL-6, and IL-12/23p40 in IFN-γ primed cells (Figures 1C and S1D). Interestingly, IL-32α did not induce production of any of the cytokines analyzed in either non-primed or primed macrophages. HSP90AA1 at 100 ng/mL induced production of TNF-α, IL-6, and IL-10 in non-primed and primed macrophages and induced IL-12/23p40 in IFN-γ primed cells. IFN-β was the only ligand with IL-10-like anti-inflammatory effects in our macrophage screens, based on the reduction in the release of TNF-α. When used at 1 ng/mL, the effect on TNF-α levels was comparable to IL-10. However, at 10-fold and 100-fold higher concentrations, IFN-β did not further reduce the levels of TNF-α (Figure 1D).

From this screening and validation work, we identified CSF-2 and IL-3 as IFN-γ-like priming ligands with respect to production of IL-6 and TNF-α but not production of IL-12/23p40 and IL-10 where different responses were observed. IL-32β was the most potent pro-inflammatory IL-32 isoform tested, triggering robust inflammatory responses in non-primed and IFN-γ primed macrophages and therefore became the focus of remaining work given the current lack of knowledge about IL-32 cytokines and their mechanism of action.

### IL-32**β** induces TLR4 and MyD88 pathway-dependent signaling in macrophages distinct from that of LPS

To date, no receptor or associated downstream signaling pathway for IL-32 cytokines has been identified and it is not clear why or how human IL-32 isoforms can elicit immunomodulatory effects in murine sytems when mice do not possess the IL-32 gene^21,22^. To gain insight into potential signaling pathways for IL-32 we stimulated various HEK293 TLR and cytokine reporter cell lines (Null, TLR4, TLR2, IFN-α/β, IFN-γ, TGF-β and TNF-α) with IL-32α, IL-32β and IL-32γ. None of the IL-32 isoforms tested induced reporter activity in HEK-Blue™ Null cells which possess a secreted embryonic alkaline phosphatase (SEAP) reporter gene under the control of the IL-12 p40 minimal promoter linked to five NF-κB/AP-1 binding sites (Figure 2A). Similarly, IL-32α, IL-32β and IL-32γ did not induce reporter activity in any of the cytokine reporter cell lines tested or in HEK-Blue™ hTLR2 cells which overexpress TLR2. However, IL-32β and, to a lesser extent IL-32γ, induced reporter activity in HEK-Blue™ hTLR4 cells which possess the same SEAP reporter gene as the HEK-Blue™ Null cells but in addition overexpress TLR4 receptor components CD14, MD-2 and TLR4 (Figure 2A). IL-32β induced NF-κB/AP-1 reporter activity in these cells with kinetics like that of LPS stimulation (Figure 2B). To validate the TLR4-dependent nature of the response to IL-32β we perturbed TLR4 signaling with TAK-242, a small molecule inhibitor of TLR4 which binds selectively to the TIR domain of TLR4 via Cys747 and blocks interactions between TLR4 and the adaptor molecules TIRAP/Mal and TRAM. TAK-242 reduced LPS-induced NF-κB/AP-1 reporter activity in HEK-Blue™ hTLR4 cells at 1 μM and IL-32β-induced activity at 0.1 μM and 1 μM (Figure 2C). Consistent with blocking effects of TAK-242, perturbation of TLR4 signaling with neutralizing antibodies to TLR4 and TLR4-MD2 also reduced LPS and IL-32β induced NF-κB/AP-1 reporter activity (Figure 2D).

**Figure 2.**
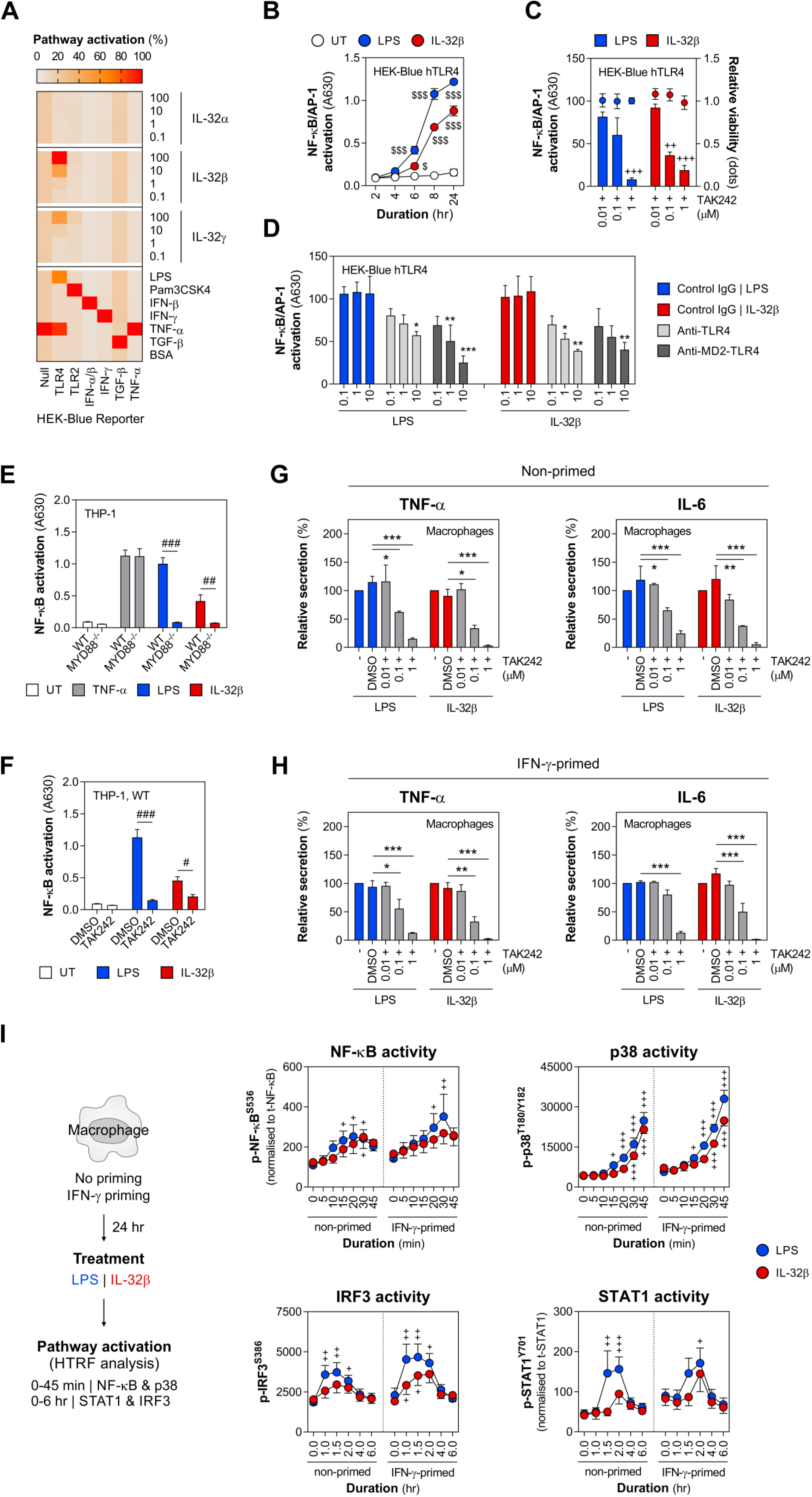
IL-32β inflammatory response is TLR4-MyD88 pathway-dependent. Also see Figure S2. (A) Heatmap showing signaling pathway activation relative to control ligands, in a panel of HEK-Blue™ reporter cell lines by three IL-32 isoforms (α, β, γ) at the indicated concentrations. (B) NF-κB/AP-1 reporter activation by LPS or IL-32β in HEK-Blue™ TLR4 reporter cells treated for the indicated times. (C) Perturbation of LPS and IL-32β-mediated NF-κB/AP-1 activation in HEK-Blue™ TLR4 reporter cells by TAK-242 and (D) TLR4 or TLR4/MD2 blocking antibodies. (E) NF-κB activation in WT and MyD88^-/-^ THP-1-Dual™ reporter cell line treated with LPS, IL-32β or TNF-α for 24 h. (F) Perturbation of NF-κB activation with TAK-242 in WT THP1-Dual™ reporter cell line. (G) Effect of TAK-242 on cytokine (TNF-α, IL-6) responses from non-primed macrophages and (H) IFN-γ-primed macrophages treated with LPS or IL-32β. (I) Experimental workflow for the analysis of signaling pathway activation in primary human macrophages by HTRF® assay at the indicated timepoints. For NF-κB and STAT1 phosphorylation normalized signal to total protein is shown. Statistical analysis: data shown in (B-D) are mean values ±SEM of 3 independent experiments with two-way ANOVA with Dunnett’s multiple comparisons test (B) vs duration-matched UT ($ p < 0.05, $$$ p < 0.001), (C) vs 0.01 TAK within LPS/IL-32β treatment groups (++ p < 0.005, +++ p < 0.001), (D) vs concentration-matched IgG control within LPS/IL-32β treatment groups (* p < 0.05, ** p < 0.005, *** p < 0.001). Data shown in (E-F) are mean values ±SEM of 4 independent experiments with two-way ANOVA with Bonferroni’s multiple comparisons test (# p < 0.05, ## p < 0.005, ### p < 0.001). Data shown in (G-H) are mean values ±SEM of n = 3 independent experiments performed on macrophages from one donor, with two-way ANOVA with Dunnett’s multiple comparisons test vs LPS/IL-32β treatment groups (* p < 0.05, ** p < 0.005, *** p < 0.001). Data shown in (I) are mean values ±SEM of n = 3 independent experiments on macrophages from one donor, with two-way ANOVA with uncorrected Fisher’s LSD test versus 0 h within LPS/IL-32β treatment groups (+ p < 0.05, ++ p < 0.005, +++ p < 0.001).

Ligand activation of TLR4 likely leads to stabilization of preformed TLR4 dimers and the recruitment of adaptors TIRAP/Mal which in turn seeds the formation of a large supramolecular organizing center (SMOC) called the Myddosome^24,25^. Myddosomes are dynamic cytosolic signaling organelle which contain the scaffolding protein MyD88 and all the adaptor, kinase and transcription factor proteins associated with and required for TLR pathway activation and inflammatory responses^26^. To investigate potential Myddosome involvement in IL-32β mediated effects in macrophages we compared LPS and IL-32β induced NF-κB activation in WT and MyD88^-/-^ KO THP-1 NF-κB reporter cell lines. Consistent with TAK-242 data from HEK-Blue™ hTLR4 cells both loss of MyD88 expression and perturbation of TLR4 signaling with TAK-242 blocked LPS and IL-32β induced NF-κB activation in THP-1 cells (Figures 2E-2F). TAK-242 also blocked LPS and IL-32β induced production of cytokines (TNF-α, IL-6, IL-23p19, IL-1β, IL-10 and IL-12p70) in non-primed and IFN-γ primed human primary macrophages (Figures 2G-2H and S2A-S2B). Due to the TLR4 pathway-dependent nature of the response to IL-32β we investigated if IL-32β itself could directly interact with components of the TLR4 complex (CD14, MD-2 and TLR4) and/or if the response to IL-32β was due to contaminating endotoxin/LPS in the IL-32β recombinant protein. To investigate a direct interaction of IL-32β with cell surface receptors, biotinylated IL-32β was added to a cell microarray of HEK293 cells overexpressing individual or combinations of TLR4 components (CD14, MD-2, TLR4) and candidate receptors for IL-32 (PR3, PAR2, αVβ3) previously reported in the literature^21^. Biotinylated LPS (LPS-EB biotin) and biotinylated anti-TGFBR2 antibody were used as positive controls. While biotinylated anti-TGFBR2 bound to cells overexpressing TGFBR2 and biotinylated LPS bound to cells overexpressing CD14 and CD14 in combination with TLR4 or/and MD-2 respectively, biotinylated IL-32β did not bind to HEK293 cells overexpressing any of the surface proteins tested (Figure S2C). To exclude the possibility of contaminating endotoxin/LPS in the recombinant IL-32β protein contributing to the TLR4-pathway dependent nature of the response to IL-32 we used the LPS neutralizing antibiotic polymyxin B to neutralize LPS in a series of comparative experiments with LPS and IL-32β in the HEK-Blue™ hTLR4 NF-κB/AP-1 reporter cell line and in human macrophages. In concentration response experiments, LPS induced NF-κB/AP-1 reporter activity in HEK-Blue™ hTLR4 cells and cytokine production from non-primed and IFN-γ primed macrophages at very low concentrations (0.1 and 1 pg/mL) compared to IL-32β (10 ng/mL) (Figure S2D). In LPS spike-in experiments in HEK-Blue™ hTLR4 cells, addition of increasing concentrations of LPS to IL-32β at 10 ng/mL did not result in additive or synergistic effects on reporter activity above that detected in response to LPS alone (Figure S2E). Polymyxin B blocked LPS induced NF-κB/AP-1 reporter activity in HEK-Blue™ hTLR4 cells and LPS-induced cytokine production in macrophages at low to moderate concentrations (0.1 pg/mL-1 ng/mL) but not fully at relatively high concentrations (10-100 ng/mL) of LPS in macrophages (Figure S2F and S2G). In comparison, polymyxin B did not block IL-32β induced NF-κB/AP-1 reporter activity in HEK-Blue™ hTLR4 cells or cytokine production in macrophages (Figure S2F and S2G). The inflammatory effects of LPS and IL-32β concentrations used in these experiments were also compared in non-primed and IFN-γ primed macrophages (Figures S2H-S2I).

Activation of canonical TLR4 pathway signaling by LPS typically leads to TIRAP/Mal-dependent seeding of Myddosome formation and possibly TRAM-dependent seeding of triffosome formation. Formation of a stable Myddosome results in activation of NF-κB, AP-1 and p38 MAPK signaling and formation of the putative triffosome leads to phosphorylation and activation of IRF3, expression of type I interferon (IFN-β) with subsequent delayed phosphorylation and activation of STAT1 due to autocrine/paracrine effects of secreted IFN-β^24,25^. To further characterize the TLR4-pathway dependent nature of the response to IL-32β we compared the kinetics and patterns of phosphorylation of NF-κB, p38MAPK, IRF3 and STAT1 in non-primed and IFN-γ primed human macrophages treated with LPS or IL-32β (Figure 2I). LPS induced significant increases in phosphorylation of NF-κB, p38MAPK, IRF3 and delayed phosphorylation – relative to IRF3 - of STAT1 in non-primed and IFN-y primed human macrophages. IL-32β also induced phosphorylation of NF-κB and p38MAPK which was similar in timing and magnitude to that of LPS but was less effective in activating IRF-3 and STAT1 in non-primed cells (Figure 2I). Taken together, these data indicate that the response of cells to IL-32β is TLR4-pathway and MyD88 dependent but is different to LPS-TLR4 canonical signaling in terms of IRF3 and STAT1 activation in non-primed macrophages.

### Identification of Myddosome components as required molecules for IL-32**β**-mediated inflammatory response in macrophages

Since the initial characterization of IL-32 as an inflammatory cytokine in 2005^27^ no cell surface signaling receptor or associated downstream signaling pathway for IL-32 has been reported in the literature^21^. To identify putative receptor/s required for cellular response to IL-32β, we performed a genome-wide RNAi screen in the HEK-Blue™ hTLR4 reporter cell line using a library of siRNA SMARTpools targeting 18,301 protein coding genes. Cells were reverse transfected with siRNAs for 72 h and then stimulated with IL-32β for 24 h to induce expression and secretion of the NF-κB/AP-1 dependent reporter gene - SEAP (Figure 3A). The genome-wide screen identified a total of 237 genes (Z-score < -2.3 and p-value < 0.01) whose perturbation led to a reduction in IL-32β-induced SEAP activity and 789 genes (Z-score > 2.5 and p-value < 0.01) whose perturbation led to an increase in IL-32β-induced SEAP activity (Figure 3B and Table S3). To identify candidate receptors and signaling pathways for IL-32β we used Metascape^28^ to perform gene annotation and biological enrichment analysis of the 237 genes whose perturbation led to a reduction in IL-32β-induced SEAP activity. Top hits were *SEC61G* and *SEC61A1* which code for the gamma and alpha subunits of the SEC61 translocon complex (Figure 3B). The SEC61 complex is located in the endoplasmic reticulum membrane and is required for secretion of soluble proteins^29^ such as the reporter SEAP therefore these hits most likely block secretion of SEAP and were not followed up in further validation work. From the list of 237 genes Metascape analysis identified the following groups of enriched GO Biological Processes (GO BPs) - MyD88-dependent toll-like receptor signaling pathway, cytokine-mediated signaling pathway, protein targeting to ER, and 3’-UTR-mediated mRNA stabilization. The genes associated with the top enriched GO BP -MyD88-dependent toll-like receptor signaling pathway - were *MyD88*, *MAP3K7/TAK1, IRAK4 and TIRAP* and additional genes required for MyD88 signaling such as *IKBKG* and *STAP2* were also hits^24,30,31^.

**Figure 3.**
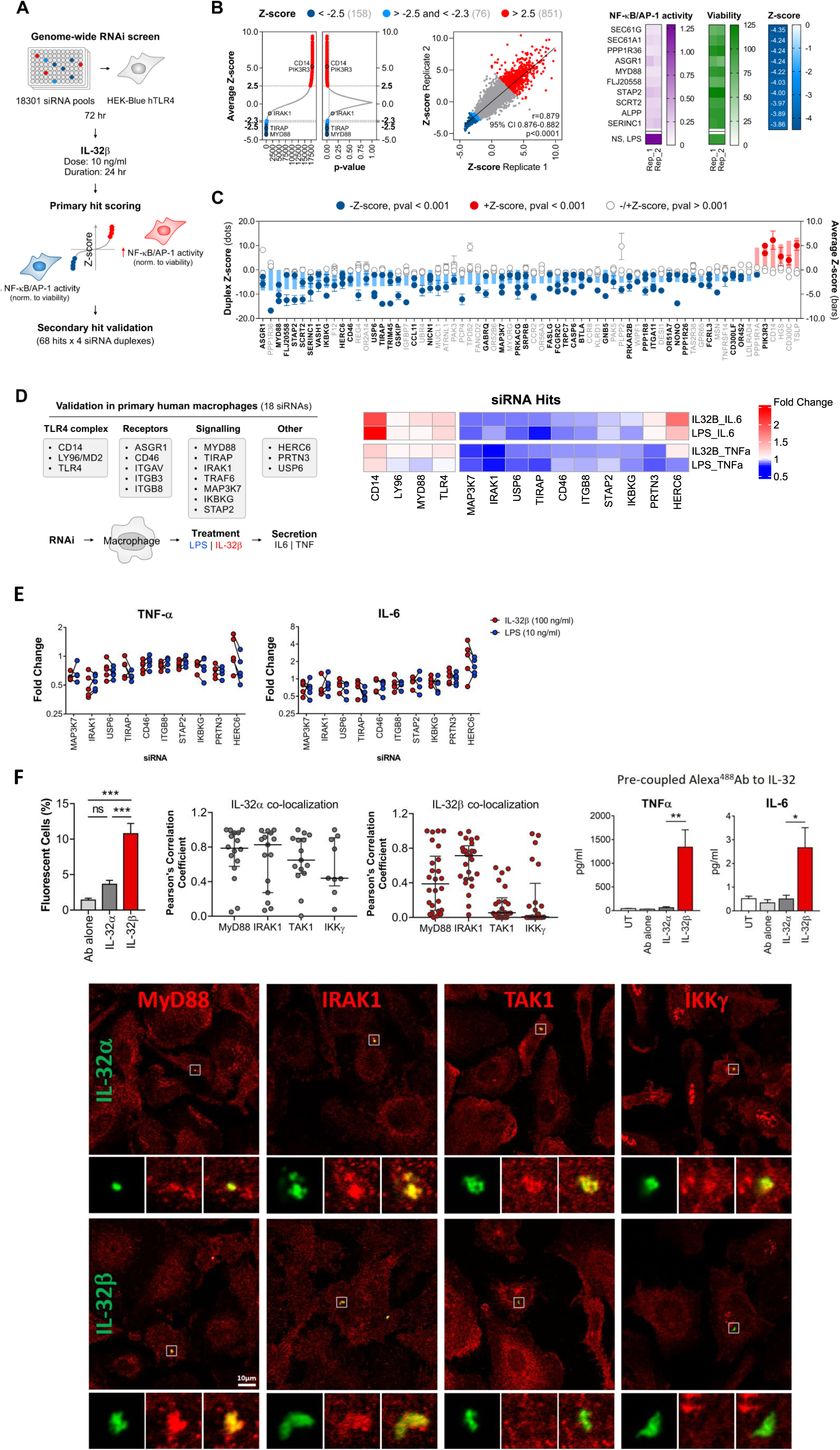
Genome-wide RNAi screen identifies Myddosome components as essential requirement for IL-32β-mediated inflammatory response in macrophages. Also see Figure S3. (A) Schematic of experimental workflow for genome-wide RNAi screen performed in HEK-Blue™ TLR4 reporter cells to identify receptors and components of the signaling pathway for IL-32β. (B) Average Z-score distribution and associated p-values across 18,301 screened genes and Z-score distribution for each of 2 SMARTpool siRNA technical replicates per gene. Heatmap of the top 10 gene hits selected based on Z-score significance is shown to the right. (C) Z-score distribution for each of 4-siRNA duplexes (dots) from deconvoluted SMARTpools for 68 selected hits in a secondary validation screen; average Z-scores from primary screen are depicted as bars. siRNA duplexes that validated in the deconvolution screen are shown highlighted in bold. (D) Schematic of experimental workflow for validation of 18 selected hits from RNAi screens in human primary macrophages (left). Heatmap showing effect of RNAi-mediated perturbation of 10 selected hits and 4 controls (CD14, LY96, MyD88, TLR4), on fold change in cytokine response from macrophages treated with LPS or IL-32β (right). (E) Variation in effect of RNAi-mediated perturbation on fold change in cytokine response after treatment with LPS or IL-32β for macrophages from individual donors. (F) Co-localization of IL-32 isoforms with hits from RNAi screens in human primary macrophages. Percentage of macrophages with a detectable fluorescent signal with IL-32α or IL-32β pre-coupled to anti-IL-32 antibody (left panel). Co-localization between IL-32 isoform-Ab complexes and Myddosome protein components was quantified with Pearson’s correlation coefficient (middle panels). Cytokine responses from human macrophages used in IF microscopy experiments to IL-32 isoform-Ab complexes (right panel). Representative images of macrophages are shown in the lower panel. Pre-coupled IL-32α (green) or IL-32β (green) co-labelled for Myddosome protein components or downstream signaling mediators (red). Statistical analysis: (F) graphs shown on the left represent mean positive cell counts from 10 random microscopy fields from 2 independent experiments. Graphs on the right represent mean cytokine release from 2 independent experiments. One-way ANOVA with Tukey’s multiple comparisons test (* p < 0.05, ** p < 0.005, *** p < 0.001) was used for both analyses.

Next, we selected genes for further validation based on Metascape gene annotation descriptors - Gene Description, Protein Function, GO BP, Subcellular Location, - in addition to manual annotation of the 237 genes prioritizing genes with supporting evidence for functions associated with receptor-mediated signaling. We compiled a list of 68 genes, which included 24 genes coding for cell surface receptors, and 6 genes whose perturbation increased IL-32β-induced SEAP activity for validation with deconvoluted SMARTpool siRNAs in the HEK-Blue™ hTLR4 reporter cell line (Table S3). The secondary deconvolution screens identified a total of 36 genes where perturbation with at least 2 of 4 siRNA duplexes (Z-score < -2.3 and p-value < 0.01) led to a reduction in IL-32β-induced SEAP activity (Figure 3C). From these 36 validated genes we prioritized and selected a total of 18 genes - including additional genes for signaling mediators in the TLR4 pathway - for RNAi mediated perturbation and validation in human primary macrophages. These genes coded for proteins involved in the TLR4 complex (*CD14, LY96/MD2, TLR4*), candidate surface receptors for IL-32β (*ASGR1, CD46, ITGAV, ITGB3, ITGB8*), proteins involved in Myddosome signaling (*MYD88, TIRAP, IRAK1, TRAF6, MAP3K7, IKBKG, STAP2*) and other additional candidate signaling proteins (*HERC6, PRTN3, USP6*) (Figure 3D). SMARTpool siRNAs targeting each of these 18 genes were transfected into primary human monocyte derived macrophages from five different donors. Macrophages were then treated with LPS (10 ng/mL) or IL-32β (100 ng/mL) for 8 h and secreted cytokines (TNF-α, IL-6) were measured as a readout (Table S4). Despite a strong donor-dependent effect on the magnitude of macrophage responses to stimuli (Figure S3A) we found 10 relevant genes (*MAP3K7/TAK1, IRAK1, USP6, TIRAP, CD46, ITGB8, STAP2, IKBKG, PRTN3, HERC6*) whose perturbation affected macrophage cytokine responses to LPS, IL-32β or both stimuli (Figure 3D and Figure S3B). Of these, MAP3K7/TAK1, IRAK1, TIRAP, STAP2 and IKBKG have all been reported to participate in TLR4-MYD88 signaling leading to activation of NF-κB^24,26^. Surprisingly, RNAi-mediated knockdown of TLR4 or MYD88 in macrophages did not significantly affect cytokine responses while *IRAK1*, which was not identified in the screen with HEK293 reporter cells, was one of the strongest hits in macrophage validation experiments (Figure 3D, Table S4 and individual values per donor in Figure 3E). Knockdown of two candidate cell surface receptors (CD46 and ITGB8) from our screens for IL-32β in macrophages elicited modest reductions in cytokine responses. While the results from this series of RNAi experiments indicated that several components of the Myddosome signaling pathway (*MYD88, TIRAP, IRAK1, MAP3K7, IKBKG, STAP2*) are required for cellular responses to IL-32β they did not definitely identify a potential receptor for this cytokine. To address this further, we selected the 24 receptor candidate genes from the primary RNAi screen data that yielded a Z-score below -2.3 and screened them for their ability to directly bind biotinylated IL-32β using Retrogenix^TM^ cell microarray technology. While the positive controls biotinylated LPS-EB and anti-TGFBR2 bound their targets CD14 and TGFBR2 respectively, direct binding of biotinylated IL-32β was not detected for any of the candidate receptors tested (Figure S3C).

To further investigate the possible association of IL-32β with proteins of the Myddosome complex, we conducted co-localization experiments using immunofluorescence confocal microscopy in primary human macrophages. To reduce background staining and increase signal specificity we pre-conjugated recombinant IL-32α or IL-32β with a fluorescently (Alexa^488^) labelled antibody which recognizes equally the main three IL-32 isoforms (α, β and γ), prior to adding these to Fcγ receptor blocked macrophages for 2h. A significantly higher number of fluorescently positive cells were identified for IL-32β than IL-32α (Figure 3F and Figure S3D). Both IL-32 isoforms displayed high levels of co-localization with MyD88 and IRAK1, measured by Pearson’s correlation coefficient (Figure 3F, middle panel). However, while IL-32α also co-localized with TAK1 and IKKγ, IL-32β did not co-localize to a significant extent with either of these proteins 2h post stimulation (Figure 3F). Consistent with previous data (Figure 1C), IL-32β but not IL-32α complexes induced significant TNF-α and IL-6 secretion from macrophages at 2 h (Figure 3F, right panel). Collectively, these results show that several components of the Myddosome signaling pathway (MYD88, TIRAP, IRAK1, MAP3K7/TAK1, IKBKG, STAP2) are required for cellular responses to IL-32β and IL-32β co-localizes with MyD88 and IRAK1 in stimulated primary human macrophages.

### IL-32**β** and LPS differ in their ability to induce immunological cross-tolerance in human macrophages

Given the fact that both IL-32β and LPS initiate signaling and inflammatory cytokine production via a Myddosome pathway in macrophages we investigated potential similarities or/and differences in their biological effects. Modeling the priming, training, and tolerance of monocytes/macrophages *in vitro* typically involves pre-treatment of cells with stimuli such as cytokines, PAMPs or DAMPs that either augment in the case of priming/training; or reduce in the case of tolerance; the induction of signature inflammatory genes in response to a secondary inflammatory stimulus^4^. These phenomena have been best characterized for LPS whereby initial pre-treatment of monocytes/macrophages with LPS can tolerize cells to subsequent treatment with LPS (tolerance) or tolerize cells to challenge with other inflammatory stimuli such as diverse PAMPs (cross-tolerance)^4^. To further investigate the effect of IL-32β on macrophages we compared the ability of IL-32β and LPS to tolerize non-primed and IFN-γ primed human macrophages to each other and to other PAMPs namely, Pam3CSK4 and zymosan. LPS tolerized non-primed and IFN-γ primed human macrophages to subsequent treatment with LPS or IL-32β, for induction of all the signature cytokines, TNF-α, IL-6 and IL-10 (Figure 4A and 4B). However, while IL-32β tolerized non-primed and IFN-γ primed macrophages to IL-32β for all these cytokines, it did not tolerize cells to LPS for induction of IL-6 and IL-12p70 by LPS (Figure 4A and 4B). Next, we investigated the ability of LPS and IL-32β to tolerize macrophages to the TLR2/TLR1 ligand Pam3CSK4 and the ability of Pam3CSK4 to tolerize cells to LPS or IL-32β. LPS and IL-32β tolerized cells to Pam3CSK4 for induction of all the cytokines measured except for IL-12p70 in IFN-γ primed human macrophages where there was slightly increased production in response to secondary stimulation with Pam3CSK4 (Figure 4C and 4D). Interestingly, Pam3CSK4 tolerized macrophages to LPS and IL-32β for induction of all the cytokines measured except for IL-6 production which was not tolerized and indeed was increased in IFN-γ primed human macrophages in response to secondary stimulation with IL-32β (Figure 4C and 4D). Finally, we investigated the ability of LPS and IL-32β to tolerize macrophages to the TLR2/TLR6 and dectin-1 ligand zymosan and the ability of zymosan to tolerize cells to LPS or IL-32β. LPS effectively tolerized cells to secondary stimulation with zymosan for induction of all the cytokines measured (Figure 4E and 4F). However, while IL-32β tolerized macrophages to zymosan for induction of TNF-α and IL-10 it did not tolerize for induction of IL-6 and IL-12p70 and it appeared to augment production of IL-6 in non-primed macrophages by zymosan. Zymosan tolerized non-primed macrophages to LPS and IL-32β for induction of the cytokines TNF-α and IL-10 but not IL-6 (Figure 4E). In IFN-γ primed macrophages, zymosan tolerized cells to LPS and IL-32β for induction of TNF-α, IL-6, and IL-12p70 production but not for induction of IL-10 where IL-10 levels were similar or greater to that measured in macrophages treated just with LPS or IL-32β (Figure 4F). In summary, it is clear from this series of tolerization experiments that IL-32β can tolerize macrophages and that IL-32β and LPS differ in their ability to induce immunological cross-tolerance both to each other and to other PAMPs. This is shown by the fact that LPS tolerized macrophages to IL-32β and zymosan for induction of all the signature cytokines, TNF-α, IL-6, IL-12p70 and IL-10 while IL-32β did not tolerize macrophages to LPS and zymosan for induction of IL-6 and IL-12p70.

**Figure 4.**
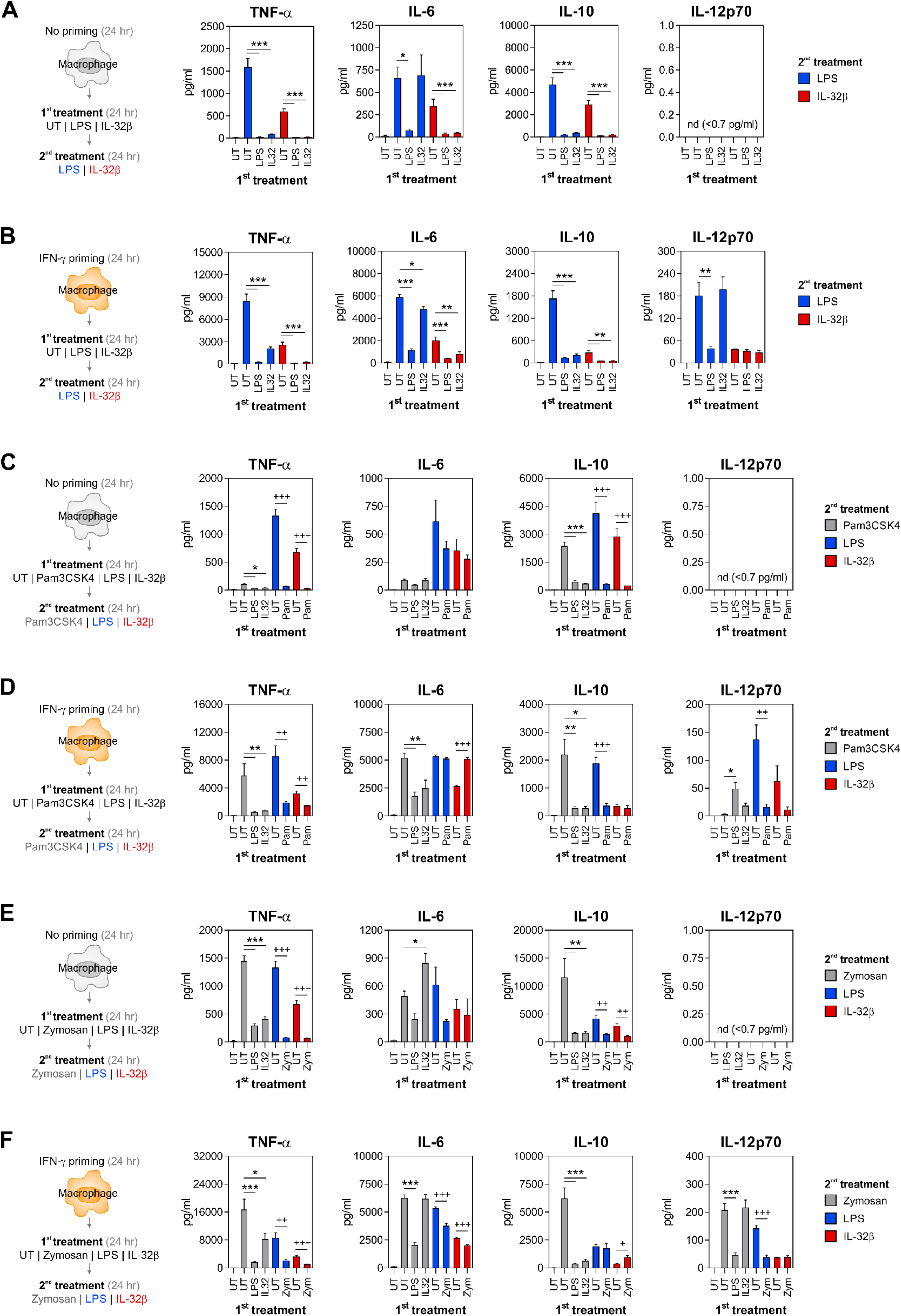
Differential induction of immunological cross-tolerance in human macrophages by LPS and IL-32β. (A-F) Schematics of experimental workflows and associated cytokine response data for induction of immunological tolerance and cross-tolerance by first (1^st^) treatment of non-primed (A, C, E) or IFN-γ-primed (B, D, F) macrophages with stimulus (LPS, IL-32β, Pam3CSK4 or zymosan) to subsequent second (2^nd^) treatment with same (tolerance) or different (cross-tolerance) stimulus (LPS, IL-32β, Pam3CSK4, zymosan). Statistical analysis: data shown in (A-F) are mean values ±SEM of n = 3 independent experiments performed on macrophages from three independent donors, with one-way ANOVA with Tukey’s multiple comparisons test (* p < 0.05, ** p < 0.005, *** p < 0.001). (C-F) For some conditions, unpaired two-tailed t-test analysis was performed as indicated (+ p < 0.05, ++ p < 0.005, +++ p < 0.001).

### IL-32**β** induces transcriptional responses in primary human macrophages which are distinct from but overlap with those induced by LPS

Our comparative analysis of macrophage responses to IL-32β and LPS identified distinct differences in terms of the activation of downstream signaling pathways (IRF-3, STAT1), and the induction of immunological tolerance and cross-tolerance. In addition, the IFN-γ-like cytokines, CSF-2 and IL-3, differed from IFN-γ with respect to their priming effects on macrophages subsequently stimulated with IL-32β and LPS. To further investigate these differences and to gain a genome-wide understanding of the effects of these cytokines and LPS on macrophage gene expression we performed transcriptomics analysis of macrophage responses to these stimuli. RNA-seq was performed on primary human monocyte-derived macrophages from n = 5 donors, primed with three priming conditions (IFN-γ, CSF2 or IFN-γ + CSF2) followed by treatment with IL-32β or LPS (Figure 5A). Analysis of cell supernatants from these experiments, confirmed stimulus induced cytokine (TNF-α, IL-6, IL-10) responses seen previously (Figure 1C and S1D) and revealed a difference in the magnitude of the effect of IL-32β and LPS on IL-12p70 and IL-23p19 production in primed cells (Figure S4A).

**Figure 5.**
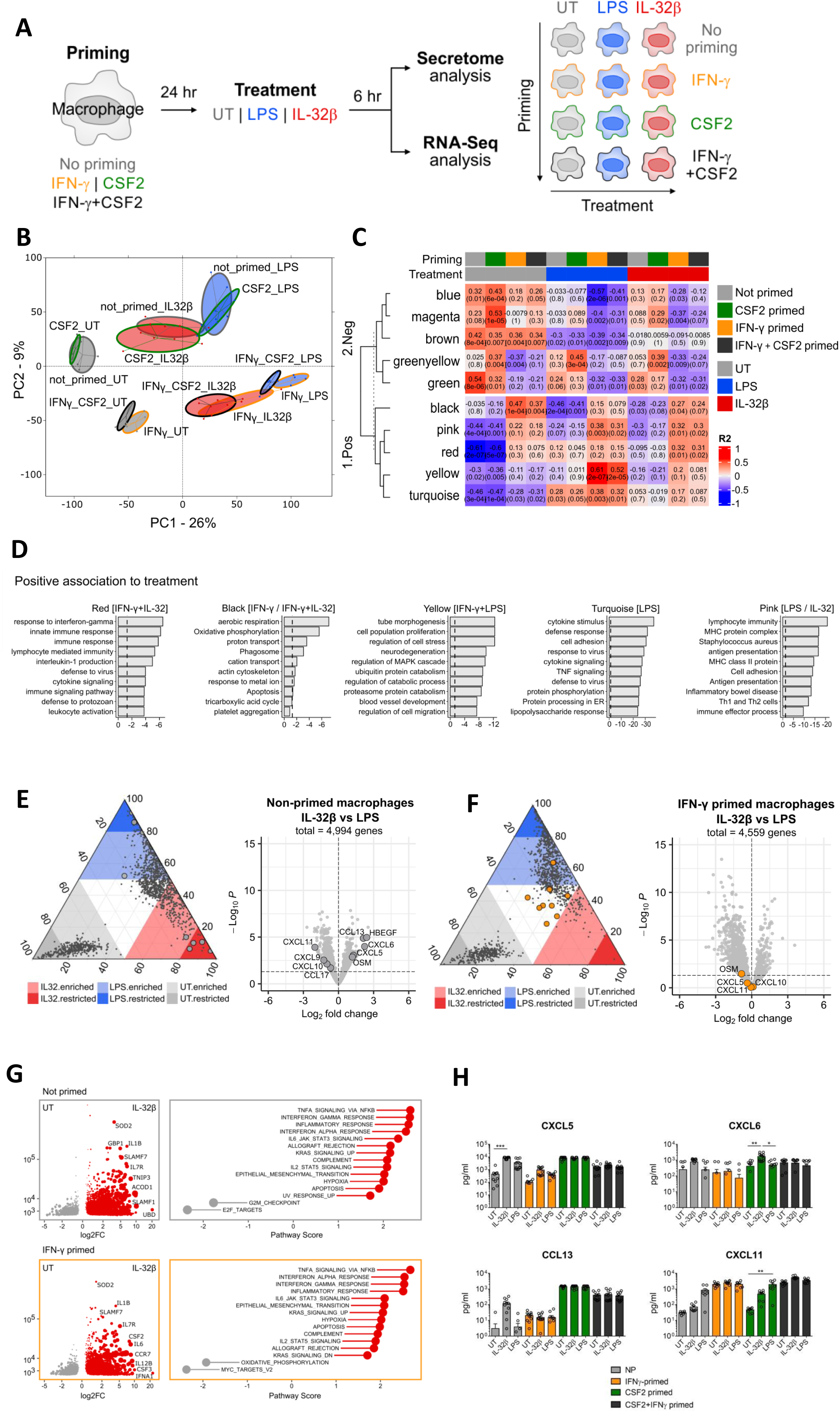
IL-32β induces gene expression changes in primary human macrophages which are distinct from but broadly overlapping with those induced by LPS. Also see Figures S4 and S5. (A) Schematic of workflow for RNA-seq experiments performed on primary human monocyte-derived macrophages from n = 5 donors, primed with three priming conditions followed by treatment with LPS or IL-32β. Treatment groups are color coded as per macrophage color scheme (schematic to right of A – outline border color of macrophage indicates priming condition, and internal color indicates treatment condition). (B) Principal Component Analysis (PCA) plot of macrophage RNA-seq gene expression profiles. (C) Module trait relationship heatmap of selected modules from WGCNA analyses that are significantly associated either positively (1.Pos) or negatively (2.Neg) to IFN-γ priming (IL32/LPS or both) treatment groups. The color scale in the heatmap (red to blue) represents the correlation between genes present in each module to the trait (treatment groups). Red color indicates positive correlation and blue indicates negative correlation. The top value inside each cell represents R^2^ value representing positive / negative correlation and the bottom value in brackets represents the p-value associated with the significance of the correlation. Modules are assigned a color identifier and row-wise clustered reflecting their positive and negative association to IFN-γ priming treatment. The top-colored bar indicates the priming status and the colored bar underneath this indicates treatment groups. (D) Barplots showing the top enriched biological terms (KEGG pathways and GO BP) for gene list in the 5 modules (*red, black, yellow, turquoise, pink*) that were positively associated to IFN-γ priming (IL32/LPS or both) treatment groups as identified by WGCNA analysis. The x-axis in the barplot represents the adjusted p-value of the enriched terms in log scale (-log10(P). The y-axis represents the selected enriched terms (shortened for aesthetic purposes). (E-F) Ternary and volcano plots comparing changes in gene expression for IL-32β and LPS treated human macrophages under non-primed (E) and IFN-γ primed (F) conditions. Genes coding for chemokines selectively induced by IL-32β or LPS in the non-primed condition are shown in both volcano plots. (G) Differentially expressed (DE) genes due to IL-32β treatment in non-primed and IFN-γ primed macrophages are shown with DESeq2 gene expression (y axis) and shrunken Log_2_FC relative to non-primed and IFN-γ primed UT conditions (x axis). Right panel indicates gene set enrichment analysis (GSEA) of these pairwise comparisons (Hallmark gene sets; MSigDB Collections). (H) Protein level validation by ELISA and MSD of chemokine genes (CXCL5, CXCL6, CCL13, CCL17, OSM, HB-EGF, CXCL9, CXCL10 and CXCL11) selectively induced by IL-32β or LPS at protein level by ELISA and MSD. Statistical analysis: data shown in (H) are mean values ±SEM of n = 5 independent experiments performed on macrophages from five independent donors in triplicates, with non-parametric Kruskal-Wallis test with Dunn’s multiple comparisons test (* p < 0.05, ** p < 0.005, *** p < 0.001).

Principal Component Analysis (PCA) of the entire RNA-seq dataset showed that treatments (UT, LPS or IL-32β) were most aligned to PC1 which accounted for 26% of the observed variance while priming conditions (IFN-γ, CSF2 or IFN-γ + CSF2) had a moderate influence on overall variance and aligned to PC2. Within the PCA, IFN-γ priming was the single most decisive factor affecting clustering of the samples. CSF2 primed samples (CSF2_UT) clustered with the non-primed sample (not_primed_UT) and IFN-γ primed samples (IFNγ_UT) co-clustered with cytokine combination (IFNγ_CSF2_UT) primed samples. This indicates that IFN-γ had stronger effects than CSF2 in eliciting transcriptional changes in macrophages in the context of priming. We next performed weighted gene co-expression network analysis (WGCNA), to determine if coordinately regulated groups of genes could be identified as overrepresented gene modules and if these modules associated with different traits i.e. treatment conditions. WGCNA identified 17 modules associated with different treatment conditions (Figure S4B). Of the 17 modules, 10 modules had a significant (p-value < 0.05) association with treatment conditions, with 5 showing negative association (blue, magenta, brown, greenyellow, green) and 5 showing positive association (black, pink, red, yellow, turquoise) (Figure 5C). Black and green modules associated with IFN-γ_priming alone; red, black and greenyellow modules associated with IFN-γ+IL32-β treatment; yellow, turquoise, blue, and brown modules associated with IFN-γ+LPS treatment and pink, magenta, and green modules associated with both combination treatment conditions (Figure 5C). Overall, IFN-γ priming strongly influenced significant module associations, which aligned with the major effect of IFN-γ on macrophage transcriptomes seen in the PCA. To gain further insight into the underlying biology we performed Metascape analysis with module genelists and present the top enriched biological terms (KEGG pathways and GO BP) for the 5 positively associated modules (Figure 5D) and 5 negatively associated modules (Figure S4D). In modules positively associated with IFN-γ and IFN-γ+IL-32 (red, black), the enriched biology included response to interferon-gamma, innate immunity, defense to virus, aerobic respiration, oxidative phosphorylation, phagosome while in the module negatively associated with these treatments (greenyellow) biology included cytoplasmic translation, ribosome and COVID-19 disease. In modules positively associated with IFN-γ+LPS (yellow, turquoise) enriched biology included tube morphogenesis, neurodegeneration, blood vessel development, response to virus, TNF signaling, lipopolysaccharide response, while in modules (blue, brown) negatively associated with this treatment, biology included ribosome biogenesis, tRNA/mRNA metabolism, HSV-1 infection, phospholipid metabolism, endosomal transport and cell projection assembly. The biology enriched in the module positively associated with both IFN-γ+IL-32 and IFN-γ+LPS (pink) included lymphocyte immunity, antigen presentation, inflammatory bowel disease while in modules negatively associated with these treatments (magenta, green) the biology included translation, DNA metabolism, mitotic cell cycle, systemic lupus erythematosus, neutrophil extracellular traps and DNA repair. The macrophage response to LPS alone was distinct from the response to IL-32β as one module (turquoise) was positively associated with LPS and IFN-γ+LPS treatment but not with IL-32β or IFN-γ+IL-32β treatment. While there were shared responses to both IL-32β and LPS in IFN-γ primed cells (pink, magenta, green modules) the response to IL-32β was characterized by biology (defense to virus, aerobic respiration, cytoplasmic translation, and COVID-19 disease) shared by IFN-γ and IFN-γ+IL-32β. Overall, this visualization and associated differential gene expression analysis indicates that IL-32β and LPS induced distinct and shared transcriptional responses in non-primed and IFN-γ primed macrophages. To gain further insight into and compare macrophage transcriptional responses to LPS and IL-32β we visualized differentially expressed genes induced by LPS and IL-32β in non-primed and IFN-γ primed macrophages using ternary plots. This analysis identified nine genes differentially regulated by both stimuli: five (*CXCL5 CXCL6 CCL13 HBEGF, OSM*) were induced exclusively by IL-32β and four (*CCL17, CXCL9, CXCL10, CXCL11*) were induced by LPS in non-primed macrophages (Figure 5E). The selectivity of these secreted ligand (chemokine, growth factor) coding genes for capturing macrophage responses to either IL-32β or LPS was abolished by priming with IFN-γ or IFN-γ+CSF2 (Figure 5F, S5A and S5B). We validated these different responses at the protein level by ELISA (Figure 5H). To further characterize the transcriptional and biological response of macrophages to stimulation with LPS or/and IL-32β, gene set enrichment analysis (GSEA) analysis was performed on lists of DEGs from non-primed and IFN-γ primed macrophages treated with IL-32β or LPS (Figure 5G and S5C). Overall, GSEA showed that the global transcriptional response to IL-32β and LPS in terms of enriched biology is largely shared. Collectively, this transcriptomics analysis indicates that IL-32β and LPS induce distinct and shared transcriptional responses in non-primed and IFN-γ primed macrophages

### IL-32 protein is elevated and an IL-32**β** gene expression signature maps to and is enriched in monocytes and macrophages in severe COVID-19

There is emerging evidence that IL-32 may be involved in several IMIDs including RA, IBD, type I diabetes (T1D) and psoriasis^21,22^. To further investigate this possibility and the potential for co-operation between IFN-γ and IL-32 in IMIDs we analyzed the expression of genes coding for *IL-32, IRAK-1, IFN-*γ and its receptors, *IFNGR1/IFNGR2* in an immune cell reference database consisting of >300,000 single-cell profiles from 125 healthy or disease-affected donors from five inflammatory diseases (RA, CD, UC, systemic lupus erythematosus (SLE), interstitial lung disease) and COVID-19 (Figure 6A-6B and Table S5)^5^. Expression of *IL-32* and *IFN-*γ genes were enriched in T cell subsets while IFN-γ receptors, *IFNGR1/IFNGR2* and the IL-32β signaling mediator *IRAK1*, were enriched in the monocyte/macrophage cell subsets in mild and severe COVID-19 (Figure 6A-6B). We also found that IL-32 protein levels were significantly higher in serum from patients with severe COVID-19 compared to mild disease and healthy controls (Figure 6C). We then generated gene expression sets or signatures for the different treatments used in our *in vitro* macrophage RNA-seq experiments (Figure 5A) and mapped these to the immune cell reference database to identify human IMID monocytes/macrophages which were transcriptionally similar. Seurat’s AddModuleScore function was used to compute an enrichment value for each gene set for each cell in the reference database and the level of enrichment was visualized along a heat color gradient on cells clustered by cell type (Figure 7). We found an enrichment of the IL-32β signature (239 genes) in the monocyte/macrophage annotated clusters isolated from patients with COVID-19 with this being most pronounced in severe COVID-19 (Figure 7A-7B, top right panel). A similar enrichment pattern was seen for gene sets from macrophages primed with IFN-γ (153 genes) or treated with IFN-γ + IL-32β (371 genes) (Figure 7B and Table S5). Interestingly, the enrichment of IFN-γ and IL-32β gene sets seemed to be selective for COVID-19 as we did not detect significant enrichment of these in the other inflammatory diseases included in the reference database. We also mapped gene sets from LPS (366 genes) and IFN-γ + LPS (527 genes) treated macrophages to the reference database and while we saw some enrichment, the magnitude was less than that observed with IL-32β (Figure S6A-S6B). Immunomodulatory drugs such as the corticosteroid, dexamethasone and the Janus kinase inhibitor (JAKinib) baricitinib or combinations of both are recommended for treatment of hospitalized patients with severe COVID-19^32^. To evaluate the effectiveness of these and other immunomodulatory drugs on blocking the inflammatory response of non-primed and IFN-γ-primed macrophages to IL-32β, we tested a panel of drugs and JAKinibs. Of the JAKinibs tested, the JAK1 selective inhibitor upadacitinib was the most effective at blocking cytokine (TNF-α, IL-6) release induced by IL-32β in IFN-γ primed macrophages (Figure 7C). Of the immunomodulatory drugs; (dexamethasone (Dx), sulfasalazine (Sf), methylprednisolone (MP)) and molecules (IL-10, anti-TNFα) tested, all except for sulfasalazine and anti-TNFα, effectively blocked cytokine (TNF, IL-6) release (Figure 7D). Taken together, these and other data support a model whereby T cell derived IFN-γ and IL-32β mediate cell-cell communication between T cells and monocytes/macrophages in COVID-19 and co-operate to drive inflammatory responses in monocytes/macrophages via JAK1/2-STAT1 and Myddosome-dependent pathways in this disease.

**Figure 6.**
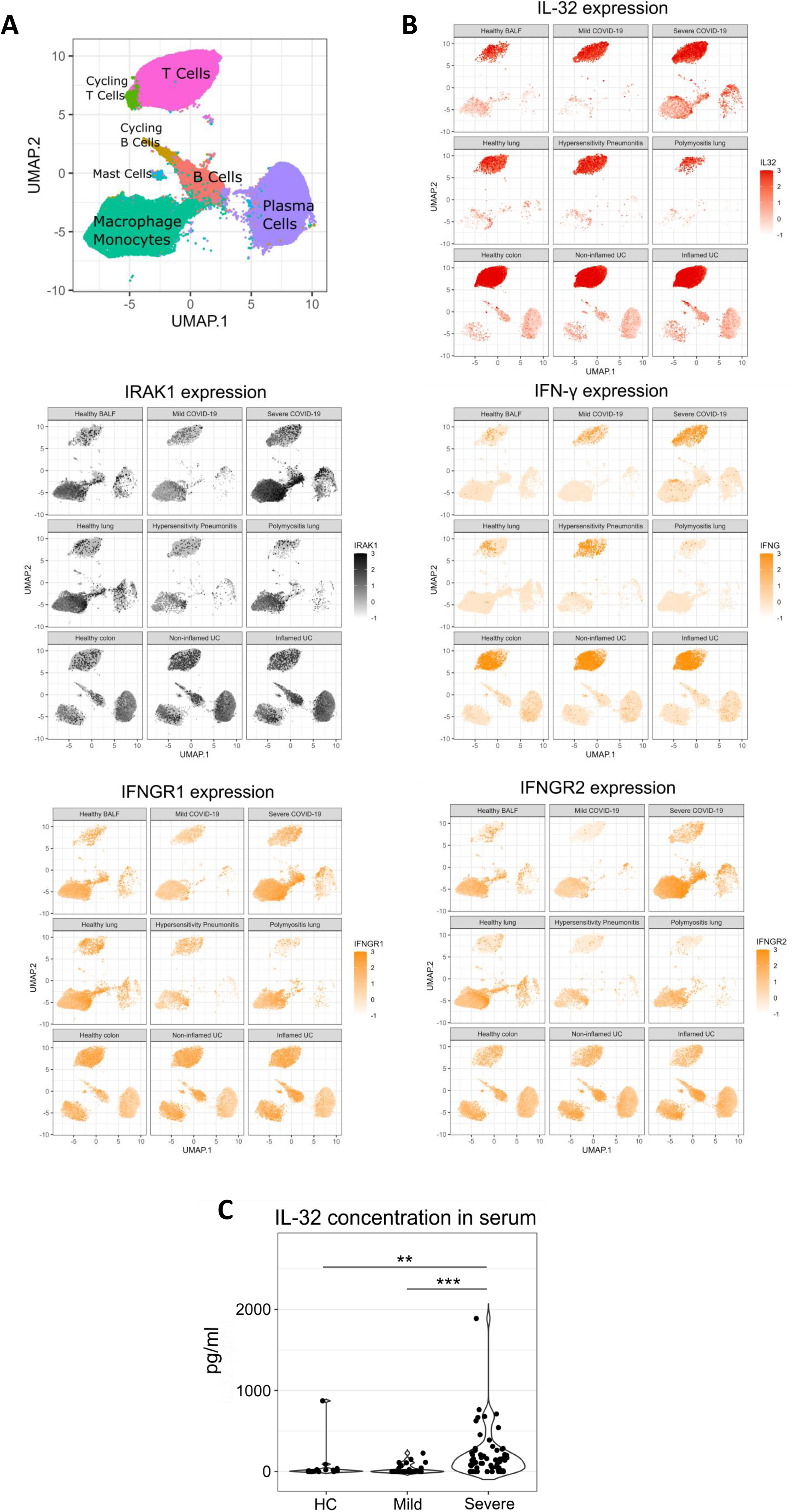
IFN-γ and IL-32 cell-cell communication pathway genes are elevated in T-cells and macrophages in COVID-19 and IL-32 protein is elevated in severe COVID-19. (A) UMAP projection schematics of the cell populations examined by single-cell sequencing ( > 300,000 single-cell profiles) from multiple inflammatory diseases and COVID-19 BALF. (B) Expression levels of the *IL32, IRAK1, IFNG, IFNGR1, IFNGR2* genes in sc-RNAseq datasets from multiple inflammatory diseases and COVID-19. (C) Detection of IL-32 protein by ELISA in serum samples from healthy donors and patients with mild and severe COVID-19. Statistical analysis: data shown in (C) are mean values of healthy individuals, HC (n = 16), mild (n = 25) and severe (n = 59) COVID-19 with non-parametric Kruskal-Wallis test with Dunn’s multiple comparisons test vs HC (** p < 0.005, *** p < 0.001).

**Figure 7.**
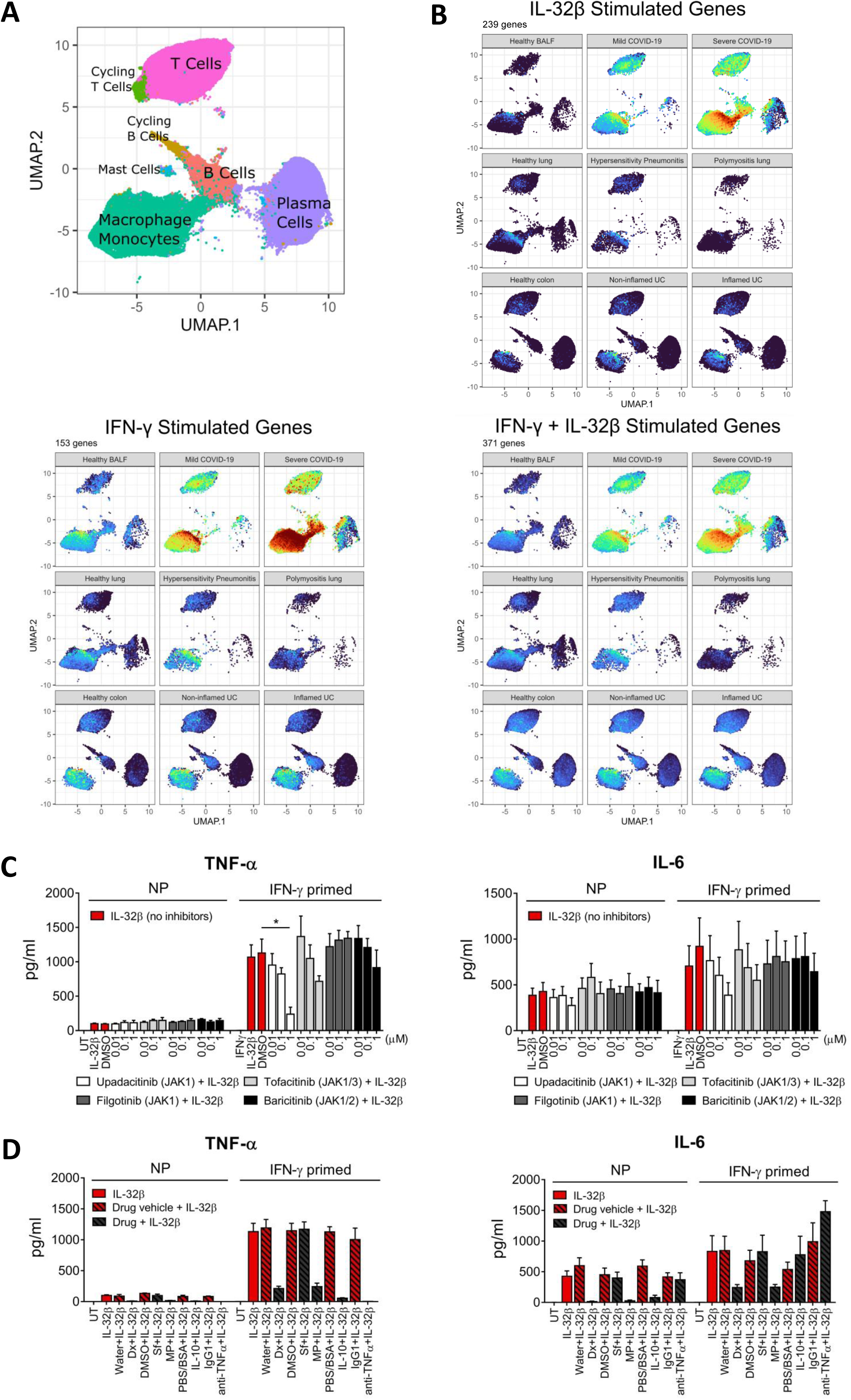
IFN-γ and IL-32β macrophage gene expression signatures are enriched in monocyte and macrophage populations in severe COVID-19. Also see Figure S6. (A) UMAP projection schematics of the cell populations examined by single-cell sequencing ( > 300,000 single-cell profiles) from multiple inflammatory diseases including COVID-19. (B) Mapping of IL-32β, IFN-γ and IFN-γ + IL-32β response gene signatures from stimulated human macrophages onto a sc-RNA sequencing atlas of multiple inflammatory diseases and COVID-19. Warmer colors indicate a higher degree of enrichment in that cell. (C) Cytokine (TNF-α, IL-6) responses from non-primed and IFN-γ primed (10 ng/mL) macrophages treated with IL-32β (10 ng/mL) +/-JAK inhibitors. (D) Cytokine (TNF-α, IL-6) responses from non-primed and IFN-γ primed (10 ng/mL) macrophages treated with IL-32β (10 ng/mL) +/-anti-inflammatory drugs. Anti-inflammatory drugs - Dx, dexamethasone; Sf, sulfasalazine; MP, methylprednisolone. Statistical analysis: data shown in (C) are mean values ±SEM of n = 3 independent experiments performed on macrophages from three independent donors with non-parametric Kruskal-Wallis test with Dunn’s multiple comparisons test as indicated (* p < 0.05).

## Discussion

IFN-γ is an essential and nonredundant cytokine for human host defense to intracellular pathogens however it is unclear if this is also the case for its role in priming M1-like macrophages and driving inflammation and immunopathogenesis in IMIDs^1,4,6,7^. In this study, we investigated if other human secreted proteins were redundant with IFN-γ in terms of its priming effects on human macrophages and/or could co-operate and synergize with IFN-γ to drive M1-like inflammatory macrophage cell states. Of 601 secreted proteins screened, we did not find any protein which could completely phenocopy IFN-γ’s transcriptional reprogramming and inflammatory priming effects on macrophages. While CSF2 and IL-3 could prime macrophages to LPS and other stimuli they did not elicit the same transcriptional reprogramming and priming effects on cells as IFN-γ. However, we did identify IL-32β, IL-32γ but not IL-32α as proteins which could co-operate with IFN-γ to drive M1-like inflammatory responses in macrophages having effects like, but distinct from those mediated by LPS. While we did not identify a receptor for IL-32β, we found that IL-32β mediated its effects on macrophages via a Myddosome dependent signaling pathway and elicited signaling, tolerance, cross-tolerance and transcriptional responses different to those induced by LPS. Analysis of IFN-γ and IL-32β gene expression and macrophage transcriptional responses to these cytokines revealed that IFN-γ and IL-32β are expressed primarily by T cells and that IFN-γ+IL-32β driven gene expression signatures are like monocyte/macrophage transcriptional phenotypes seen in mild and severe COVID-19. Supporting these observations we also found that IL-32 protein was elevated in serum from patients with severe COVID-19. Overall, our data supports (i) a signaling model where host T cell-derived IL-32β can signal to macrophages via a Myddosome-dependent pathway eliciting effects similar to, yet distinct from those induced by microbial derived LPS and (ii) an immunopathogenic model where T cell secreted IFN-γ and IL-32β mediate cell-cell communication between T cells and macrophages and co-operate to drive inflammatory macrophage responses via JAK1/2-STAT1 and Myddosome-dependent signaling pathways in severe COVID-19.

IL-32 was originally reported in 1992 as a human gene named NK4 which encoded a 27kD protein expressed selectively in human lymphocytes^33^. Expression of the gene was increased after activation of T cells with mitogens and NK cells by IL-2. The gene structure, splice variants, regulation and pro-inflammatory effects of the protein on THP-1 cells and murine RAW 264.7 macrophages was first reported in 2005^27^. Because of its ability to induce inflammatory cytokines (TNF-α, IL-8, MIP-2) in human and murine cells and to activate the NF-κB and p38MAPK signaling pathways in murine macrophages it was identified as a cytokine and renamed IL-32^27^. Despite its cytokine-like biology, the IL-32 gene has no sequence homology with any other cytokine families, is found only in humans with eight orthologues in primates and no other orthologues found in any of 239 other species including rodents. The IL-32 gene has 35 transcript variants (ensemble.org) of which 30 potentially encode proteins^22^. Of these, nine protein isoforms have been validated experimentally and four isoforms IL-32α, IL-32β, IL-32γ and IL-32ε have been characterized most extensively in the literature both in human and murine model systems^21,22^. The IL-32γ isoform was previously reported to be the most active isoform but IL-32β had equivalent biological activity to IL-32γ − with IL-32α having minimal activity - in PBMC assays reported in that study^23^. Over the past 20 years it has become clear that IL-32 is expressed constitutively by many human cell types including immune (NK cells, T cells), non-immune (epithelial, endothelial, fibroblasts) and cancer cells. *IL-32* expression can also be induced and increased in response to various types of bacterial and viral infections and in response to cell type selective inflammatory stimuli such as IL-18, IL-12, IFN-γ, TNF-α, IL-1β, anti-CD3/CD28, ConA, LPS^21,22^. While IL-32 isoforms lack a signal peptide for processing through the ER-Golgi secretory pathway they appear to be secreted from cells via different unconventional protein secretion (UPS) mechanisms which may be context, isoform, cell type and mode of release dependent. Indeed, IL-32 protein is detected in cell culture supernatants of various stimulated cell types (epithelial cells, T cells) and is elevated in cells, serum and tissues from several IMIDs such as, T1D, SLE, RA and IBD^34–37^. In addition, several viral and bacterial infections increase IL-32 expression and both intracellular and secreted extracellular IL-32 isoforms have anti-microbial effects, for example IL-32γ has been shown to inhibit influenza virus, HIV and HBV replication^21,22,38^. Amongst immune cells, the highest expression is found in T cell subsets (memory CD4^+^ Treg, CD4^+^ Th1) and recently IL-32β was identified as the predominant IL-32 isoform secreted by human T cells as a free protein via a UPS mechanism in response to IL-2 and TCR stimulation^22,39^. One of the enduring unanswered questions about IL-32 biology concerns the identity of a receptor and downstream signaling pathway for IL-32 isoforms in target cells (monocytes, macrophages, fibroblasts, T cells) which could explain their largely pro-inflammatory effects and association with IMID pathology. This question is perplexing because even though the IL-32 gene is only found in humans and primates human IL-32 isoforms stimulate inflammatory responses from murine macrophages and elicit inflammatory, anti-inflammatory, tolerogenic and disease modifying effects in different IL-32 transgenic (TG) mice^21,22^.

In this study, IL-32γ was identified as ligand with M1-like inflammatory activity in macrophage immunoinflammatory screens of 601 IMID-relevant recombinant proteins. Subsequently, IL-32γ and IL-32β but not IL-32α were validated as ligands which could induce inflammatory responses in non-primed and primed (IFN-γ, CSF-2, IL-3) macrophages with IL-32β eliciting the most potent effects. To the best of our knowledge, this is the first report of IL-32 isoforms co-operating and synergizing with IFN-γ to stimulate inflammatory cytokine responses from human macrophages and is like previous reports where IFN-γ has been shown to co-operate and synergise with LPS or TNF-α^3,5^. Given the current lack of knowledge about IL-32 cytokine biology in terms of known receptors and downstream signaling pathways we investigated potential mechanisms underpinning its effects on human cells and macrophages using complementary experimental approaches. As an initial approach, we screened a panel of HEK293 TLR and cytokine reporter cell lines (Null, TLR4, TLR2, IFN-α/β, IFN-γ, TGF-β and TNF-α) for responsiveness to IL-32 isoforms and found that HEK-Blue™ hTLR4 cells which overexpress TLR4 receptor components (CD14, MD-2 and TLR4) responded robustly to IL-32β. The CD14/MD-2/TLR4 complex recognizes, binds to and is activated by LPS and has also been reported to recognize other PAMPs and proteins such as heat shock proteins (HSPs) and amyloid--beta (Aβ) aggregates^24,40^. To exclude the potential contribution of contaminating LPS to observed IL-32β responses we used the LPS neutralizing antibiotic polymyxin B to neutralize LPS in a series of concentration response experiments in HEK-Blue™ hTLR4 cells and human macrophages and found that polymyxin B did not block IL-32β responses. In addition, biotinylated IL-32β did not bind to the CD14/MD-2/TLR4 complex or any previously identified candidate receptors (PR3, PAR2, αVβ3) for IL-32β in a cell-based microarray. To validate the TLR4 pathway-dependent nature of the response to IL-32β we perturbed the pathway in HEK-Blue™ hTLR4 cells, THP-1 NF-κB reporter cells and human macrophages with the TLR4 antagonist TAK242^41^ and showed that it blocked responses to LPS and IL-32β. In addition, MyD88^-/-^ KO THP-1 NF-κB reporter cells did not respond to LPS or IL-32β. While these data showed that the response of human cells to IL-32β was TLR4 and MyD88 pathway-dependent and independent of contaminating LPS or direct binding of IL-32β to the CD14/MD-2/TLR4 complex they did not identify the receptor/s. The TLR4-dependent nature of the response is like previous observations in murine peritoneal macrophages from *Tlr4* wt C57BL/6 and *Tlr4* mt C3H/HeNj mice which have a single point mutation in the Toll/IL-1R domain of TLR4. In that study the cytokine response of macrophages from LPS-resistant *Tlr4* mt C3H/HeNj mice to a panel of IL-32 isoforms including IL-32β was much less than the response from *Tlr4* wt cells^23^. We reasoned that the overexpression of CD14/MD-2/TLR4 in the HEK-Blue™ hTLR4 cell line somehow made them competent to respond to IL-32β and therefore performed an unbiased genome-wide RNAi screen in these cells to identify the potential receptor/s and additional signaling pathway components for this unconventional cytokine. The RNAi screens in HEK-Blue™ hTLR4 cells and RNAi validation in primary human macrophages identified several components of the Myddosome signaling pathway (*MYD88, TIRAP, IRAK1, MAP3K7, IKBKG, STAP2*) as been required for cellular responses to IL-32β. Despite the identification of 24 candidate receptors for IL-32β from these screens none of them validated convincingly in RNAi experiments, neutralization experiments (ASGR1, CD46 – data not shown) nor did biotinylated IL-32β bind to any of them in a cell based microarray. To gain further insight into this biology we investigated the possible association of IL-32β and IL-32α with proteins of the Myddosome complex and found that IL-32α co-localized with MyD88, IRAK TAK1 and IKKγ while IL-32β co-localized with MyD88 and IRAK1 in stimulated primary human macrophages. A possible explanation for the low number of large fluorescently labelled punctate structures observed in these experiments could be due to the formation of supramolecular organizing centers (SMOCs). Stimulus triggered oligomerization and assembly of signaling proteins such as MyD88 into SMOCs called Myddosomes is a well-known phenomenon for receptors involved in innate immune responses like TLR4 and IL-1Rs^42^. Mechanistic insights from TLR4, IL1R1 and Myddosome signaling studies indicate that Myddosome size and stability can act as a threshold which determines whether signaling is propagated in the cell^25,26,43,44^. The large punctate staining patterns observed for IL-32β and co-localized Myddosome proteins are like those typically seen when MyD88 oligomerizes into Myddosome SMOCs^43^. It is possible, that overexpression of CD14/MD-2/TLR4 in the HEK-Blue™ hTLR4 cell line promotes oligomerization and assembly of MyD88 into Myddosomes, priming cells for activation and downstream Myddosome signaling via either direct or indirect interaction with IL-32β but this remains to be formally investigated. Collectively these data indicate that the response of human cells and macrophages to IL-32β is MyD88 and Myddosome dependent and perhaps receptor independent. It will be interesting to see if IL-32β directly interacts with components of the Myddosome and if this Myddosome dependency is also the case for IL-32γ and other IL-32 isoforms which have similar biological activity. A prediction based on these findings is that murine *Myd88* KO macrophages and IL-32 TG mice on a *Myd88* KO background would be largely resistant to the inflammatory and disease modifying effects of human IL-32β and perhaps other IL32 isoforms.

Given the fact that both IL-32β and LPS initiated signaling and inflammatory cytokine production via a Myddosome pathway we compared their effects on non-primed and IFN-γ primed macrophages to identify potential similarities or/and differences in their overall biological effects. Consistent with the literature we found that both LPS and IL-32β activated the NF-κB and p38MAPK signaling pathways in non-primed and IFN-γ primed macrophages however while LPS activated IRF3 and STAT1 in non-primed macrophages IL-32β did not. This indicates that IL-32β, unlike LPS, does not effectively activate TRAM-dependent seeding of the triffosome leading to activation of IRF3, expression of type I interferon (IFN-β) and delayed phosphorylation and activation of STAT1 due to the autocrine/paracrine effects of secreted IFN-β^25^. This difference in signaling between LPS and IL-32β in terms of IRF3 and STAT1 activation in non-primed macrophages was consistent with differences in transcriptional responses from these cells as detected by RNA-seq analysis. Overall, we found that while relatively few DEGs distinguished LPS and IL-32β responses, LPS but not IL-32β was able to increase expression of the interferon stimulated and STAT1 regulated chemokine genes CXCL9, CXCL10 and CXCL11 which aligned with activation of IRF3 and STAT1 by LPS but not IL-32β. We also found that LPS and IL-32β differed in their ability to induce immunological tolerance and cross-tolerance in macrophages. We report for the first time that IL-32β tolerized macrophages to itself and that IL-32β and LPS differed in their ability to induce immunological cross-tolerance both to each other and to other PAMPs. While LPS tolerized macrophages to IL-32β and zymosan for induction of all signature cytokines, TNF-α, IL-6, IL-12p70 and IL-10 analyzed, IL-32β did not tolerize macrophages to LPS and zymosan for induction of IL-6 and IL-12p70. These findings are consistent with and may provide a mechanistic explanation for previous divergent observations in various IL-32 TG mice where human IL-32β and IL-32γ appeared to elicit inflammatory, anti-inflammatory and tolerogenic effects depending on the model used^21^. For example, IL-32γ TG mice were reported to be more resistant to LPS induced endotoxin shock and expression of IL-32β in IL-32β TG mice was reported to reduce inflammatory cytokines and disease pathology in a model of arthritis^45,46^. Again, a prediction of these findings is that these divergent effects of human IL32 isoforms in TG mice would be absent on a *Myd88* KO background. Another implication of these data is that host derived IL-32β could elicit host protective anti-inflammatory tolerogenic responses to itself but not to microbial derived PAMPs such as LPS and zymosan. This could conceivably be an important difference for maintaining a robust innate immune response in the context of infectious disease and sepsis. Our global transcriptional analysis of non-primed and primed (IFN-γ, CSF2 or IFN-γ + CSF2) macrophage responses to LPS and IL-32β further confirmed the non-redundant nature of the priming and transcriptional reprogramming effects of IFN-γ on macrophages compared to CSF2. Overall, IFN-γ priming strongly influenced significant module associations to LPS and IL-32β seen in WGCNA and this aligned with the major effect of IFN-γ on macrophage transcriptomes seen in the PCA. This analysis also showed that IL-32β and LPS induced distinct and shared transcriptional responses in non-primed and IFN-γ primed macrophages and that enriched biology such as defense to virus, oxidative phosphorylation, phagosome and COVID-19 disease were shared in IFN-γ and IFN-γ+IL-32 treated macrophages.

Recently it has become clear that an IFN inflammatory signature in macrophages (and other cell types) is a common shared transcriptional phenotype and cell state associated with multiple IMIDs, infectious diseases such COVID-19 and inadequate therapeutic responses to anti-TNFs^5,11,13–19^. In addition, there is evidence that IL-32 may be involved in several IMIDs including RA, IBD, T1D, psoriasis and in bacterial and viral infectious diseases^21,22,38^. To gain further insight into the potential roles of both IFN-γ and IL-32β in human disease we analyzed the expression of genes coding for the IL-32 and IFN-γ pathways and gene expression signatures from macrophages treated with IL-32β, IFN-γ or IFN-γ+IL-32β in an immune single cell reference database from five IMIDs and COVID-19^5^. We found that expression of *IFN-γ* and *IL32* ligand genes were enriched in T cells while expression of IFN-γ receptor and Myddosome component *IRAK1* were enriched in monocyte/macrophage cells in COVID-19. In addition, IFN-γ, IL-32β and IFN-γ+IL-32β induced response signature genes were enriched in monocyte/macrophage annotated clusters isolated from patients with COVID-19 with this being most pronounced in severe COVID-19. In support of these findings IL-32 protein levels were significantly higher in serum from patients with severe COVID-19. Furthermore, we found that dexamethasone which is recommended for treatment of hospitalized patients with severe COVID-19^32^ and elicits its therapeutic effects by suppressing pathological interferon responses in monocytes^47^ effectively blocked cytokine release induced by IL-32β in IFN-γ primed macrophages. Taken together, these and other data support a model whereby by T cell derived IFN-γ and IL-32β mediate cell-cell communication between T cells and macrophages and co-operate to drive inflammatory responses in macrophages via JAK1/2-STAT1 and Myddosome-dependent pathways in COVID-19. This model is consistent with previous work reporting existence of an IFN-γ driven positive feedback loop between T cells and inflammatory monocyte/macrophage populations and the overall contribution of monocytes and macrophages to pathological inflammation and cytokine storm in severe COVID-19^5,14,48–53^. It will be interesting to see if this model can be validated in murine and human co-culture systems and to what extent this co-operation between IFN-γ and IL-32 contributes to immune responses and immunopathology in other human IMIDs and infectious diseases.

## Limitations of the study

Collectively, our data shows that the unconventional cytokine, IL-32β activates non-primed and IFN-γ primed human macrophages via a Myddosome signaling pathway and has effects on macrophages which are distinct yet overlapping with those triggered by LPS through TLR4 mediated activation of the canonical Myddosome pathway. However, we could not identify a cell surface receptor for IL-32β despite the use of several complementary experimental approaches such as genome-wide RNAi screens, RNAi validation screens in macrophages, flow cytometry assays and cell-based microarray receptor screens with biotinylated IL-32β. It is possible that other approaches such as biased or genome-wide CRISPR/Cas9 knock-out screens in appropriate target cell types, may identify a receptor for IL-32 isoforms, but this remains to be seen. Therefore, it remains an open question as to how IL-32β and other IL-32 isoforms initiate signaling on target cells. What is clear from our work is that IL-32α and IL-32β can enter cells and co-localize with Myddosome components but how this happens and whether these IL-32 isoforms directly interact with MyD88, IRAK1, TAK1 and IKKγ is unknown at this time.

Finally, while we demonstrated that gene expression signatures from IFN-γ, IL-32β and IFN-γ+IL-32β treated macrophages were enriched in monocyte/macrophage clusters in severe COVID-19 this association may not be causal.

## Resource availability Lead contact

Further information and requests for resources and reagents may be directed to, and will be fulfilled by, the lead contact Ken Nally.

## Materials availability

This work did not generate new unique reagents and components.

## Data and code availability

The data discussed in this publication have been deposited in NCBI’s Gene Expression Omnibus and are accessible through GEO Series accession number GSE287623 (https://www.ncbi.nlm.nih.gov/geo/query/acc.cgi?acc= GSE287623)

## Acknowledgments

This work was supported by grants from Science Foundation Ireland—namely a career development award (CDA) to K.N. (SFI-13/CDA/2171), a research center grant (SFI-12/RC/2273) and a research center academic-industry partner Spoke award (SFI-14/SP/2710) to K.N, F.S. and APC Microbiome Ireland.

## Author Contributions

A.R-V. and A.S. performed experiments and analysed data. A.S. drafted figures. A.R-V. drafted figures and the paper. T.C. performed informatics analysis, vizualisations of RNA-seq data, analysed single-cell sequencing data and provided technical and conceptual help and J.V performed the cell-based screen data analysis, processed the sequencing data through bioinformatics pipeline, performed WGCNA, enrichment analysis (metascape) and relevant RNA-seq visualizations. C.L. performed the RNAi validation work and provided technical and conceptual help. A.J.L.(2) performed the microscopy experiments. J.W. analyzed data, drafted figures, formatted all figures and performed statistical analysis. P.S. performed RNA library preparation. W.C.A. and L.O.M. recruited subjects and provided serum from healthy controls and COVID-19 patients. I.B.J provided conceptual and oversight guidance and advice on statistical and bioinformatics analysis. A.J.L.(1) provided project management and technical advice. S.M. provided technical and conceptual help. O.N. performed biotinylation of IL-32β. R.S. provided technical and conceptual help with proteomic work. M.Macoritto and S.C. provided help with data curation and target validation. M.C.L. B.L.McR., M.Matzelle and K.N. conceived the study and provided conceptual help. M.Matzelle provided technical advice, coordinated experimental work. K.N. and F.S. obtained grant funding. K.N. conceived and designed all experiments, supervised the research and wrote the final manuscript. All authors reviewed data and manuscript.

## STAR Methods

### EXPERIMENTAL MODEL AND SUBJECT DETAILS

#### Human monocyte-derived macrophages

Cryopreserved human monocytes from healthy donors isolated by CD14 positive selection were purchased from BioIVT. Monocytes were seeded at a density of 0.5 x 10^6^ cells/mL (1 x 10^5^ cells per well) in 96-well plates (Nunc™ Edge 2.0, Nunclon™Δ treated) in RPMI-1640 medium, supplemented with 10% (v/v) heat-inactivated fetal bovine serum (Merck-Sigma, F9665), 100 U/mL penicillin, 100 μg/mL streptomycin and 100 ng/mL recombinant human M-CSF (Peprotech, 300-25) and cultured for five days to differentiate monocytes into macrophages. M-CSF was added fresh on day 5 by replacing the culture media and including or not the cytokine priming treatment. To prime macrophages, IFN-γ (10 or 25 ng/mL) was added on day 5 for 24 h and media was then aspirated from cells after this 24h priming, cells were then washed with sterile PBS prior to addition of subsequent stimuli. For macrophage based phenotypic screens and tolerance experiments IFN-γ was used at 25 ng/mL, for all other experiments, IFN-γ was used at 10 ng/mL.

#### HEK293 and THP1 reporter cell lines

Commercial HEK-Blue™ reporter cell lines stably transfected with plasmids bearing human TLR genes or receptors for human cytokines were purchased from InvivoGen (TLR4, hkb-htlr4; TLR2, hkb-htlr2; parental cell line, hkb-null2; IFN-α/β, hkb-ifnab; IFN-γ, hkb-ifng; TGF-β, hkb-tgfb; TNF-α, hkb-tnfdmyd). Cells were maintained in DMEM (ThermoFisher, 31966021) 4.5 g/liter glucose, 2 mM L-glutamine, supplemented with 10% (v/v) heat-inactivated fetal bovine serum, 100 U/mL penicillin, 100 μg/mL streptomycin, 100 μg/mL Normocin™ (InvivoGen, ant-nr-1) and selection antibiotic formulations from InvivoGen, following manufacture’s guidelines. To avoid potential cell loss during treatment, 96-well culture plates were prepared before the cell seeding by pre-coating wells with 200 μl of poly-L-lysine (Sigma, P4707) for 24 h at 4°C and then wells were subsequently washed with sterile PBS. Cells were seeded at a density of 7.5 x 10^4^ cells/mL and treated with the indicated stimuli for 24 h. Pathway-dependent secreted embryonic alkaline phosphatase (SEAP) activation was determined using QUANTI-Blue™ solution (InvivoGen, rep-qbs), according to manufacturer’s instructions. Optical density at 630 nm was measured using BioTek™ Synergy 2 microplate reader. Wild-type (WT) and MyD88 knockout (MyD8^-/-^) THP1-Dual™ reporter cell lines were purchased from InvivoGen (InvivoGen, THP1-Dual™ Cells, thpd-nfis and THP1-Dual™ KO-MyD thpd-komyd). THP1 cell lines were cultured in RPMI-1640 medium, 2 mM L-glutamine, 25 mM HEPES, supplemented with 10% (v/v) heat inactivated fetal bovine serum, 100 U/mL penicillin, 100 μg/mL streptomycin, 100 μg/mL Normocin™ and selection antibiotics Zeocin® and blasticidin, following manufacturer’s guidelines. Cells were seeded at a density of 1 x 10^5^ cells/mL 24 h prior to treatment for additional 24 h. SEAP activation was determined using QUANTI-Blue™ solution. Recombinant human TNF-α (Peprotech, 300-01A) at 25 ng/mL was used as a positive control for WT and MyD88^-/-^ cell NF-κB reporter activation.

### METHOD DETAILS

#### Gene list of 1,516 Crohn’s disease relevant ligand-receptor genes

To select a human immune-mediated inflammatory disease (IMID) relevant gene list and library of secreted ligand proteins for macrophage-based phenotypic screens we performed a comprehensive bioinformatics analysis of six publicly available Crohn’s disease datasets^54–58^. To filter for relevant genes, we sourced additional gene level information from the human protein database (http://www.proteinatlas.org/) and searching for gene ontology terms that correspond to secretome/receptome networks in the immune system: extracellular region (GO.0005576), extracellular space (GO.0005615), extracellular exosome (GO.0070062), cytosol (GO.0005829), integral component of membrane (GO.0016021), cell surface (GO.0009986), cytoplasm (GO.0005737) and integral component of plasma membrane (GO.0005887). Upstream regulators inferred from ingenuity pathway analysis performed on gene lists from Arijs et al., 2009^56^, Haberman et al., 2014^57^ and annotated secreted and signaling receptome proteins^59^ were also selected and integrated. Genes were given a scoring based on 3 criteria: gene expression in the diseased tissue, gene function and ligand or receptor score and subsequently ranked. After applying these filters 1,516 genes of interest were selected. Our gene list was further validated using an additional RNA-seq dataset of biobanked paired non-inflamed and inflamed colonic mucosal biopsies from patients with active CD (n = 47) and control group (n = 28)^60^. Based on this bioinformatic analysis, we assembled a list of 601 proteins and peptides that were sourced commercially for macrophage-based phenotypic screens (Table S1).

#### Human macrophage immunoinflammatory screens

Cell-based phenotypic screens in human macrophages were performed using a library of 601 human recombinant protein ligands (582 proteins and 19 peptides). Cell culture grade bioactive recombinant human proteins were purchased from Biotechne-R&D Systems^TM^ and peptides were purchased from Tocris. Proteins and peptides were reconstituted in sterile PBS with 0.1% BSA. The stock protein ligand library was arrayed in eight 96-well polypropylene plates (Thermo Scientific™, 10644502) at a 100-fold concentration (recombinant proteins at 10 µg/mL and peptides at 100μM) in sterile PBS with 0.1% BSA. For primary screens, a 10-fold dilution of the master stock plates was prepared in PBS/0.1% BSA in eight daughter plates. The final concentration of recombinant proteins in test plates with macrophages was 100 ng/mL and for peptides was 1 μM. Parameters of the screens were optimized in a series of pilot experiments testing optimal monocyte/macrophage cell density for differentiation and treatments, duration of treatments, concentrations of proteins and selection of secreted cytokine readouts (data not shown). To analyze the biological response of macrophages in response to the ligand library, we measured the release of cytokines (TNF-α, IL-6, IL-12/IL-23p40, IL-10, IL-27) which are used to evaluate inflammatory, priming, innate immune training, and tolerance responses in macrophages. All chosen cytokines are elevated and involved in immunopathogenesis of different immune-mediated inflammatory diseases (IMIDs). For all screens, monocyte-derived human macrophages from one blood donor were used in triplicate. Macrophages were exposed to the ligand library under four different experimental workflows. Workflows were designed to identify ligands with effects on macrophages similar to that of (i) LPS (LPS-like or pro-inflammatory activity), (ii) IFN-γ (IFN-γ-like priming effect to LPS or IFN-γ-like activity), (iii) IFN-γ + LPS (LPS-like effect in IFN-γ-primed cells or M1-like activity), and (iv) IL-10 (IL-10-like effect or anti-inflammatory activity). Pro-inflammatory activity, macrophages were left unprimed and treated on day 6 of differentiation with the ligand library for 24 h; IFN-γ-like activity, macrophages were primed with the ligand library for 24h on day 5, then washed with sterile PBS and treated with LPS for 24 h on day 6; M1-like activity (polarization to M1 macrophage inflammatory phenotype in IFN-γ-primed cells), macrophages were primed with IFN-γ for 24h on day 5 and after washing with sterile PBS, cells were immediately treated with the ligand library for 24 h on day 6; anti-inflammatory activity, macrophages were treated with the ligand library for 6 h on day 5 and primed with IFN-γ on day 5, followed by treatment on day 6 with LPS for 24 h. Untreated macrophages and macrophages primed with IFN-γ were included in each test plate. The following controls were used as reference for cell response to stimuli: pro-inflammatory activity, treatment with LPS at 50 ng/mL for 24 h; M1-like and IFN-γ-like priming activity, priming with IFN-γ at 25 ng/mL and subsequent treatment with LPS for 24 h; anti-inflammatory activity, treatment with IL-10 at 100 ng/mL for 6h, priming with IFN-γ for 24 h and treatment with LPS for 24 h. On day 7, cell supernatants were collected to determine release of the cytokines: TNF-α, IL-6, IL-12/IL-23p40, IL-27 and IL-10 were analyzed by MSD assay. Relative cell viability was determined by CellTiter-Glo Assay, CTG (Promega, G7570), following manufacturer’s instructions. Secondary validation screens were performed on a selection of hits following the same experimental procedure as described for primary screens. The activity of selected hits from the secondary screen was further evaluated by means of concentration response experiments, treating differentiated macrophages with 10-fold increasing concentrations of protein ligands, ranging from 0.1 to 100 ng/mL.

#### Analysis of human macrophage immunoinflammatory screen data

A data analysis method based on altered Z-scores was used due to its strength in identifying outliers (i.e ligands that produce a response which deviate from the normally distributed background). Data analysis was performed in three steps: (1) data normalization, (2) p-and q-value calculations and (3) data filtering. Data from eight test plates was normalized using altered Z-score median centered normalization method. Next, the p- and q-values were calculated to associate the altered Z-scores to statistical significance and allow ranking of the proteins. Using the altered Z-scores, the p-values were calculated using pnorm function and two-tailed test in R. The p-values were then converted to q-values estimating false discovery rate (FDR) using R package (qvalue) with the default settings. Finally, filters were applied to the data based on Z scores and q-values. For pro-inflammatory, IFN-γ-like and M1-like activity, ligands were selected as hits if above filter threshold: (+) Z4 + qvalue (0.05). For, anti-inflammatory, negative Z score (-) Z4 + qvalue (0.05) was used. After applying these filters, we obtained the following the number of hits: 42 (pro-inflammatory), 43 (M1-like), 9 (IFN-γ-like) and 34 (anti-inflammatory). An extended subset of 82 proteins per condition were manually selected for secondary screens. The secondary screen data was analysed by calculating the median values of released cytokines from three independent replicates per condition. These values were relativized to the median value of the positive control, LPS, and expressed as a percentage. The hits with the highest values were selected for further validation using concentration response experiments:

#### HEK293 reporter cell line assays

The following HEK293 reporter cell lines were used to investigate responses to IL-32 proteins. HEK-Blue™ Null2 cells (hkb-null2), selection reagent Zeocin® (ant-zn); HEK-Blue™ hTLR4 (hkb-htlr4) and HEK-Blue™ hTLR2 (hkb-htlr2), selection reagent HEK-Blue™ Selection (hb-sel); IFN-α/β reporter HEK293 cells (hkb-ifnab) and IFN-γ reporter HEK293 cells (hkb-ifng), selection reagents Zeocin® and blasticidin (ant-bl) respectively; TGF-β reporter HEK293 cells (hkb-tgfbv2), and TNF-α reporter HEK 293 cells (hkb-tnfdmyd), selection reagents Zeocin® and puromycin (ant-pr) respectively. The following positive control ligands were used to verify reporter cell activation: LPS-EK Ultrapure (InvivoGen, tlrl-peklps) at 10 ng/mL, Pam3CSK4 (InvivoGen, tlrl-pms) at 1 μg/mL, recombinant human IFN-β (Biotechne-RnD, 8499-IF) at 1 ng/mL, recombinant human IFN-γ (Biotechne-RnD, 285-IF-100/CF) at 1 ng/mL, recombinant human TGF-β (Biotechne-RnD, TGFB2-015H) at 1 ng/mL, recombinant human TNF-α (Promega, 300-01A) at 1 ng/mL. For pathway activation kinetics, cells were treated with LPS (10 ng/mL) or IL-32β (100 ng/mL) for the indicated times.

#### Perturbation of TLR4 pathway activation

Neutralizing monoclonal antibodies targeting human TLR4 (InvivoGen, mabg-htlr4) or TLR4/MD2 (InvivoGen, mab-htlr4md) receptors and IgG1 isotype control (InvivoGen, mabg1-ctrlm) were used on HEK-Blue™ hTLR4 cells at the concentrations: 0.1-1-10 μg/mL. Cells were pre-treated for 1 h with neutralizing antibodies before treating cells with LPS-EK Ultrapure at 1 ng/mL or IL-32β (Biotechne-RnD, 6769-IL-025) at 10 ng/mL for 24 h. The TLR4-specific small molecule inhibitor TAK-242 (Cayman, 13871) was used at concentrations: 0.01-0.1-1 μM on HEK-Blue™ cells and non-primed or IFN-γ-primed human macrophages. Macrophages were pre-treated with TAK-242 for 1 h at 37°C before treating cells with LPS-EK Ultrapure at 1 ng/mL, LPS from *E. coli* O111:B4 (Merck-Sigma, L3024 or IL-32β at 10 ng/mL for 24 h. In THP1-Dual™ cells, TAK-242 was used at a single concentration of 1 μM and cells were pre-treated with TAK-242 for 1 h before treating cells with LPS-EK Ultrapure at 10 ng/mL and IL-32β at 100 ng/mL for 24 h.

#### Endotoxin/LPS spike-in experiments

For endotoxin/LPS spike-in experiments, HEK-Blue™ hTLR4 cells and differentiated human macrophages were treated with a concentration series of 10-fold dilutions of LPS (0.1 pg/mL to 100 ng/mL) or IL-32β (0.01-100 ng/mL). Activation of TLR4-dependent NF-κB/AP-1 reporter activity in HEK-Blue™ hTLR4 cells and release of TNF-α and IL-6 from macrophages was analysed by QUANTI-Blue™ assay or MSD, respectively. To quantify the effect of increasing concentrations of spike-in endotoxin/LPS on IL-32β protein triggered TLR4-dependent NF-κB/AP-1 reporter activity, 10-fold dilutions of LPS (0.1 pg/mL to 100 ng/mL) were added to a single concentration of IL-32β (10 ng/mL). Cells were incubated for 24 h and TLR4 activation was measured by QUANTI-Blue™ assay.

#### Neutralization of endotoxin/LPS with polymyxin B

Polymyxin B (InvivoGen, tlrl-pmb) was prepared in sterile water according to manufactureŕs instructions. A concentration series of 10-fold dilutions of LPS (0.1 pg/mL to 100 ng/mL) and IL-32β (0.01-100 ng/mL) was mixed with polymyxin B (10 μg/mL) or culture media and incubated at 37°C for 2 h. These treatments were then added to HEK-Blue™ hTLR4 cells or macrophages from n = 3 independent donors. HEK-Blue™ hTLR4 and macrophage supernatants were analysed 24 h later with QUANTI-Blue™ assay or MSD assay for TNF-α and IL-6 respectively.

#### IL-32**β** Biotinylation

Recombinant Human IL-32β [R&D Systems, 6769-IL] in 20 mM HEPES, 500 mM NaCl, 1 mM DTT, and 0.2% (w/v) CHAPS, pH 7.0, was labeled with Sulfo ChromaLINK® Biotin [Vector Lab, B-1007] at a 75X molar ratio for 90 minutes at room temperature to overcome reduced reactivity due to the presence of CHAPS and DTT. The sample was quenched with Tris, pH 7.0, and excess biotin was removed by dialysis using a Slide-A-Lyzer G2 Dialysis Cassette, 3.5K MWCO [Thermo Scientific, 87725], followed by one round of Zeba Spin Desalting Columns, 40K MWCO [Thermo Scientific, A57766]. Protein concentration, degree of labeling, and size heterogeneity were assessed by UV/Vis absorbance and size exclusion chromatography.

#### Retrogenix^R^ cell microarray receptor screens

To identify candidate IL-32β surface receptor(s) we utilized the human cell microarray technology screening service provided by Retrogenix^TM^. Expression vectors carrying reporter gene ZsGreen1 and selected candidate human receptors were spotted on a microarray slide in duplicates. Individual vectors or combinations of vectors used were: TLR4 (gene ID: 7099), MD2 (gene ID: 23643), CD14 (gene ID: 929), TLR4 + MD2, TLR4 + CD14, MD2 + CD14, TLR4 + MD2 + CD14, ITGAV (gene ID: 3685), ITGB3 (gene ID: 3690), ITGAV + ITGB3, PRTN3 (gene ID: 5657), F2RL1 (gene ID: 2150, mRNA version BC018130), F2RL1 (gene ID: 2150, mRNA version NM_005242), ITGB6 (gene ID: 3694), ITGAV + ITGB6. Additionally, TGFBR2 (gene ID: 7048), and EGFR (gene ID: 1956) were included as positive controls. Human HEK293 cells were reverse transfected with the spotted vectors and cultured to allow receptors to be overexpressed. Biotinylated IL-32β was added at 10 μg/mL for 5 min at 37°C in DMEM with 10% FBS before slide fixation. Biotinylated anti-TGFBR2 antibody at 1 µg/mL was used as a positive control. Optimization of binding conditions to TLR4 receptor complex was performed with LPS-EB Biotin (InvivoGen, tlrl-lpsbiot) at 2 μg/mL. Binding to candidate receptor proteins was assessed by fluorescent imaging using Alexa^647^ streptavidin. Slide images were analysed using ImageQuant software (GE Healthcare Life Sciences). Binding test for LPS and IL-32 to the 24 receptors identified in the GW siRNA secondary deconvolution screen was performed by reverse transfection of human expression vectors for the proteins: ASGR1 (gene ID: 432), REG4 (gene ID: 83998), OR2A14 (gene ID: 135941), ATRNL1 (gene ID: 26033), GABRQ (gene ID: 55879), OR52B6 (gene ID: 340980), SRPRB (gene ID: 58477), OR56A3 (gene ID: 390083), CCR2 (gene ID: 729230), FCGR2C (gene ID: 9103), TRPC7 (gene ID: 57113), CCR8 (gene ID: 1237), BTLA (gene ID: 151888), KLRD1 (gene ID: 3824), ITGA11 (gene ID: 22801), OR51A7 (gene ID: 119687), TAS2R38 (gene ID: 5726), GPR65 (gene ID: 8477), FCRH3 (gene ID: 115352), MSN (gene ID: 4478), TNFRSF14 (gene ID: 8764), NKIR (gene ID: 146722), OR4S2 (gene ID: 219431), C18ORF1 (gene ID: 753), 3 isoforms of CD46 (gene ID: 4179). The following 5 controls were included based on bibliographic references and our preliminary results: ASGR2 (gene ID: 433), TLR4 (gene ID: 7099), CD14 (gene ID: 929), MD2 (gene ID: 23643), PRTN3 (gene ID: 5657), in addition to 2 internal controls TGFBR2 (gene ID: 7048), and EGFR (gene ID: 1956), were used bringing to 34 the final number of candidate receptors screend. HEK293 cells were fixed and incubated with 1 µg/mL of IL-32β, 2 µg/mL of LPS-EB Biotin or 1 µg/mL of biotinylated anti-TGFBR2 antibody or PBS for 30 mins at 37°C in DMEM with 10% FBS.

#### Homogeneous Time Resolved Fluorescence (HTRF®) assays

Human monocyte-derived macrophages seeded at a density of 0.5 x 10^6^ cells/mL in 96-well plates were left either non-primed or primed with IFN-γ for 24 h before treating them with LPS or IL-32β (both at 10 ng/mL) in duplicate over a time course,. The selected timepoints for analyzing each phospho-protein of interest were: 0, 5, 10, 15, 20, 30 and 45 minutes for pNFKB and p38; 0, 1, 1.5, 2, 4 and 6 hours for pIRF3 and pSTAT1. Detection of phosphorylation on IRF3 (Ser386), p38 (Thr180/Tyr182), NFKB (Ser536) and STAT1 (Tyr701) proteins was performed with homogeneous time-resolved fluorescence (HTRF®) kits (Cisbio, 6FRF3PEG, 64P38PEG, 64NFBPEG and 63ADK026PEG, respectively) according to manufactureŕs instructions. Briefly, cells were lysed with 50 μL of supplemented lysis buffer and 16 μL of the lysate was used for the assay, in duplicates. Measurements were made with BioTek™ Synergy 2 microplate reader after 24 h of incubation with d2-Eu Cryptate antibody mix. Fluorescent signal for phospho-NFKB and phospho-STAT1 was normalized to total NFKB and total STAT1 protein signal with HTRF® kits (Cisbio, 64NFTPEG and 63ADK096PEG, respectively) following the kit indications: Normalization value = (Phospho HTRF Ratio/Total HTRF Ratio) x 100. Not normalized values for IRF3 and p38 are represented as HTRF Ratio = (Signal 665 nm / Signal 620 nm) x 10^4^.

#### Human macrophage tolerization experiments

Induction of immunological tolerance and cross-tolerance in non-primed and IFN-γ primed human macrophages by LPS, IL-32β, TLR2/TLR1 ligand Pam3CSK4 (InvivoGen, tlrl-pms) and TLR2/dectin1 ligand zymosan (InvivoGen, tlrl-zyn) was investigated in human macrophages. Cells were left non-primed or primed with IFN-γ or for 24 h, incubated with LPS or IL-32β at 10 ng/mL, Pam3CSK4 at 1 μg/mL or zymosan at 5 μg/mL for 24 h and challenged again for additional 24 h. Supernatants were collected and analysed by MSD or LEGENDplex^TM^ assay.

#### Genome-wide RNAi screens

Genome-wide RNAi screens to identify genes required for TLR4-dependent NF-κB/AP-1 reporter activity in HEK-Blue™ hTLR4 cells in response to IL-32β treatment was performed by the functional genomics core at SBPMDI (Sanford Burnham Prebys Medical Discovery Institute, La Jolla, CA, USA). The screen was run twice in 384-well format using the Dharmacon genome-wide human ON-TARGET-Plus (OTP) siRNA library, containing a single SMARTpool of 4 siRNAs (10 nM) targeting 18,301 genes. Two assays were used as screen readouts: TLR4-dependent-NF-κB/AP-1 SEAP reporter activity was determined using QUANTI-Blue™ and cell viability as an internal normalization control was determined by CellTiter-Glo® (CTG) Assay. A large batch of low-passage HEK-Blue™ hTLR4 reporter cells sufficient for the entire screening process were prepared and frozen together. Cells were maintained in DMEM with 10% of FBS, supplemented with 100 μg/mL Normocin and selection antibiotics following supplier’s instructions. Antibiotics were removed during transfection and screening process. Parameters of the screen were optimized in a series of pilot experiments testing optimal cell density, efficiency, and toxicity of a panel of transfection reagents, concentration of IL-32β and duration of the treatment. In the optimization process, positive control target genes were chosen from the canonical TLR4 pathway across a range of expected phenotypic strength: TLR4, MYD88, CD14, AP-1 and NFKB and cells were treated with the titration of IL-32β, LPS and TNF-α. For the final screen, cells were seeded at a density of 2,000 cells/well, transfected with siRNA (10 nM) using Lipofectamine RNAiMAX for 72 h (Invitrogen; Thermo Fisher Scientific, Inc., 13778075) before the IL-32β treatment at 10 ng/mL for 24 h. Negative controls included transfection lipid alone and two kinds of non-specific siRNAs: 4 mixed siRNAs, that control for the assay and 48 mixed siRNAs, used for checking screen background and normalization. To analyse primary siRNA screen data: first, SEAP values were normalized with CTG values, then, the median of the sample population in the plate and median absolute deviation (MAD) were used to calculate the robust Z-score, following the formula: z* = (xij – median(x))/MAD * 1.4826. A secondary deconvolution screen was performed on selected 68 gene hits in duplicates, based on normalized average Z-score from primary screen and bibliographic references. To do so, the 4 individual siRNA duplexes that comprise the gene specific SMARTpools were transfected into the HEK-Blue™ TLR4 reporter cell line for the corresponding selected targets. The complete SMARTpools for the gene hits were used as control for the secondary deconvoluted screen.

#### Macrophage RNAi validation screens

RNAi validation of 14 genes identified from the genome wide RNAi screen and 4 comparative control genes in the TLR4 pathway (CD14, LY96, MyD88, TLR4) was performed in human macrophages using siRNA SMARTpools (ON-TARGETplus technology, Horizon Discovery). 1 pmol of SMARTpool siRNA was prepared in OptiMEM (Thermo Scientific™, 31985047) and mixed with Lipofectamine RNAiMAX (Thermo Scientific™, 13778075) according to manufacturer’s instructions. Reverse transfection was performed by plating 10 μl siRNA-lipid complex per well in 96-well plates. Macrophages were harvested on day 5 with Accutase™ (Corning, 15323609), diluted to a density of 1 x 10^6^ cells/mL and 90 μl of cells were distributed on the siRNA-lipid complexes. After 24 h 100 μL of complete media was added to each well. After 48 h cells were washed and cultured in 200 μl of complete media. Macrophages were treated 72 h post-transfection with 10 ng/mL of LPS or 100 ng/mL of IL32β for 8 h.

#### Cytokine and chemokine assays

Cytokine analysis in all macrophage supernatants was performed by ELISA, MSD electrochemiluminescence assay or LEGENDplex^TM^ assay, following manufacturer’s instructions. DuoSet ELISA kits from Biotechne-RnD were used to detect CXCL5 (DY254), CXCL6 (DY333), CCL13 (DY327), CCL17 (DY364), OSM (DY295), HB-EGF (DY259B) and IL-32 (DY3040). Cytokines TNF-α, IL-6, IL-12/IL-23p40, IL-27, IL-10 and chemokines CXCL9, CXCL10 and CXCL11 were analyzed by electrochemiluminescence (MSD, Meso Scale Diagnostics). Detection of IL-12p70, IL-23p19, IL-1β and IL-10 in supernatants from macrophage RNA-seq experiment, macrophage TAK242 and tolerance experiments was performed with LEGENDplex^TM^ assay in V-bottom plates and analysed with BD Celesta flow cytometer. Acquired data was analysed with LEGENDplex^TM^ Data Analysis Software. The supernatants collected at 6 h from RNA-seq experiments were also used to analyse the IL-32 or LPS selective release of chemokines. Supernatants from n = 4 different donors were diluted with RPMI media as indicated and analysed for the following chemokines in a final volume of 50μL: CXCL5 (1:9), CXCL6 (1:3), CCL13 (1:3), CCL17 (1:1), OSM (1:3) and HB-EGF (no dilution).

Supernatants from n = 3 different donors diluted at 1:1 ratio were used for analysing CXCL9, CXCL10 and CXCL11. Analysis of IL-32 in serum samples from COVID-19 patients^61^ was performed by diluting samples with 10% FBS in PBS to a final volume of 50 μL. Serum from healthy individuals (n = 16), pre-screened for any acute or chronic immune, metabolic or infectious disorder, was obtained in Cork, Ireland before COVID-19 pandemic outbreak. Serum was obtained from hospitalized patients with mild COVID-19 from Ticino, Switzerland (n = 25) or with severe COVID-19 from St. Gallen, Switzerland (n = 23) and Geneva, Switzerland (n = 36).

#### IL-32 co-localization, immunofluorescence and confocal microscopy

Human recombinant IL-32α or IL-32β were prepared (2 μg/mL) in sterile PBS with 0.5% BSA in microcentrifuge tubes. For microscopy, each IL-32 protein was mixed with 2.5 μg of Alexa^488^conjugated anti-IL32 antibody (Biotechne-RnD, IC30402G). Equal volumes of IL-32 proteins and antibody were mixed and incubated for 30 min on ice in the dark to allow binding before adding to the cells. Human monocytes (0.25 x 10^6^ cells) were seeded on 13 mm sterile glass coverslips placed in 24-well plates and differentiated for 5 days with M-CSF (100 ng/mL). Macrophage Fcγ receptors were blocked using 12.5 μg per 10^6^ cells of Human BD Fc Block™ (BD Biosciences, 564219) in RPMI for 15 min at RT before the addition of IL-32^AF488^ pre-conjugated IL-32 protein antibody complexes. Macrophages were incubated with IL-32^AF488^ for 2 h at 37°C in the dark. Coverslips were washed with warm PBS and cells were fixed with 4% paraformaldehyde in PBS for 15 minutes at RT and quenched with 50 mM NH_4_Cl in PBS for 15 minutes at RT, followed by blocking and permeabilization with 0.05% saponin/0.2% BSA in PBS for 30 minutes at RT. The coverslips were then transferred to a humid chamber and incubated with primary antibodies for detection of protein components in the Myddosome: MyD88 (Thermo Fisher, PA5-19919), IRAK1 (CST, 4359), IKKγ/NEMO (Thermo Fisher, MA5-32682) and TAK1/MAP3K7 (Thermo Fisher, 700113) in blocking solution for 1 h at RT. A secondary Cy3-conjugated anti-rabbit antibody (Jackson ImmunoResearch, 711-165-152) was then used for detection. Nuclei were labelled with DAPI and the coverslips were mounted on glass slides with MOWIOL. To quantify the percentage of cells with IL-32-positive structures at least 10 random fields were chosen per condition, from two biological replicates performed in duplicate, under the DAPI filter. The number of cells displaying Alexa^488^–positive intracellular structures were counted per field. Over 400 cells were quantified per condition. Images were acquired with a Zeiss LSM 510 laser scanning confocal microscope (Carl Zeiss, Jena). For the colocalization studies, at least 5 random fields were chosen per condition from three biological replicates and images were acquired using a 63x/1.4 numerical aperture Plan Apo objective lens. Pearson’s colocalization coefficient was calculated using Zeiss ZEN software.

#### Treatment of macrophages with anti-inflammatory drugs

The following JAK inhibitors were used: upadacitinib (Insight Biotechnology, HY-19569), tofacitinib (Cayman Chemical, CP-690550), filgotinib (Selleckchem, S7605) and baricitinib (Selleckchem, S2851) at concentrations of 10 mM, 100 mM and 1 μM in DMSO. The following anti-inflammatory drugs were also used at the indicated concentrations: dexamethasone (Sigma, D2915) 100 nM in water, sulfasalazine (Sigma, BP779) 3 μM in DMSO, methylprednisolone (Cayman Chemical, 15013) 200 nM in DMSO, IL-10 (Biotechne-RnD, 217-IL-005/CF) 100 ng/mL in 0.1% BSA in PBS, anti-TNFα (Research Grade Adalimumab biosimilar, Biotechne-RnD, MAB9677) 1 μg/mL and IgG1 Isotype Control (MSC, 403502) 1 μg/mL in PBS. Macrophages were pre-treated with JAK inhibitors and drugs for 30 minutes before priming with IFN-γ. On the following day macrophages were washed with sterile PBS and macrophages were again pre-treated with JAK inhibitors and drugs for 30 minutes before finally treating the cells with LPS or IL-32β (10 ng/mL).

#### Macrophage RNA-seq

RNA-seq was performed on primary human monocyte-derived macrophages from n = 5 donors, primed with three priming conditions followed by treatment with LPS or IL-32β. Macrophages were left non-primed, or primed with IFN-γ, CSF2 or IFN-γ + CSF2 (10 ng/mL each) for 24h and then treated with LPS or IL-32β (10 ng/mL) for 6 h. RNA was isolated using RNeasy Micro Kit (QIAGEN, 74004) according to manufacturer’s instructions. Cells were lysed in 350 μl RLT buffer supplemented with β-mercaptoethanol and RNA was isolated using the RNeasy Micro Kit and eluted from columns in nuclease-free water. RNA concentration was determined using Qubit RNA Broad-Range Assay Kit (Invitrogen, Q1021) and quality of purified RNA was determined with Agilent 2200 TapeStation system. All samples yielded an RNA integrity number (RIN) above 8.5. RNA-seq library was generated with RNA 6000 PICO Kit (Agilent, 5067-1513). A total amount of 20 μg RNA per sample was used as input material for the library preparations. Library size was determined by Agilent 2200 TapeStation system and library concentration was quantified by PCR using Kappa Library Quantification Kit (Roche, 07960298001). All libraries were pooled at an equimolar concentration. Sequencing was carried out by Source BioScience, using a NovaSeq 6000 platform (Illumina) on a 150bp paired-end run. The 60 RNA-seq libraries were sequenced for paired-end reads. The FASTQ files from the 60 samples were processed through the analysis pipeline. On average, 44.5 million reads (one direction) were obtained for each sample with an average Phred score of 35. The read quality was examined using FastQC (Babraham Bioinformatics) and low-quality reads were trimmed using Fastp^62^. The trimmed reads were pseudo aligned to the human transcriptome (hg38, gencode_v36) using kallisto (with options -b 100, -genomebam -gtf)^63^. The R package tximport^64^ was used to aggregate the transcript per million counts (TPM) quantified at the transcript level counts to gene level counts. Normalization method of trimmed mean of M-values (TMM)^65^ was performed using the calcNormFactors function in edgeR^66^. On average 69% of reads were aligned to human transcriptome. In total there were 19,671 protein coding genes and 39,756 non-protein coding genes. We focused rest of our analyses on the protein coding genes. R function filterByExpr was used to determine which genes had sufficiently large counts to be retained in a statistical analysis. In total, 15,586 out of 19,671 protein coding genes were retained after using this function.

#### Principal Component Analysis

Normalized log (cpm) voom counts of 15,586 genes were used as an input for principal component analysis (PCA) using R function dudi.pca from ade4 R package^67^. The first five principal components accounted for 26%, 9%, 7%, 5% and 5% variance respectively. Priming status and treatment labels were combined and used as a grouping factor. The fill color in eclipse represents the treatment and the border color the priming status. Function s.class from ade4graphics R package was used to represent a two-dimensional scatter plot grouping points along axes-1 and axes-2.

#### Weighted Gene Co-expression Network Analysis (WGCNA)

The WGCNA package^68^ in R was used to perform a weighted gene co-expression network analysis^69^ to cluster the 15,586 genes into modules. The workflow for the WGCNA analysis comprised the following steps: 1. automatic network construction and module detection were performed using WGCNA blockwiseModules function (power=14, corType = “pearson”, networkType = “signed”, TOMType = “signed”, mergeCutHeight=0.15, minModuleSize=100, depSplit=4). This involved constructing an adjacency matrix, computing into a Topological Overlap Matrix (TOM), module identification using merged dynamic and dynamic tree cut algorithm. In total, 17 modules were identified at this setting. 2. Correlations among gene expression modules and phenotypic traits were investigated and significance was determined by a Student asymptotic p-value for the given correlations. The modules were associated with the following traits - not_primed_UT, CSF2_primed_UT, IFN-g_primed_UT, IFN-g_CSF2_primed_UT, not_primed_IL32b, CSF2_primed_IL32b, IFN-g_primed_IL32b, IFNg_CSF2_primed_IL32b, not_primed_LPS, CSF2_primed_LPS, IFN-g_primed_LPS IFNg_CSF2_primed_LPS. Of the 17 modules, 10 modules that had a significant association (p-value < 0.05) to the IFNg_priming condition with either IL32, LPS or both were selected for further investigation. Of the 10 modules that had significant association to IFN-γ priming, 5 showed positive association (*pink, red, black, yellow, turquoise*) and 5 showed negative association (*blue, magenta, brown, greenyellow, green*).

#### Differential expression analysis, gene set enrichment analysis (GSEA) ternary, volcano and upset plots generation

Transcript abundance files were imported into the Deseq2 analysis pipeline using the tximport package. Normalization was performed using the Deseq function and results generated using the Results function with logFC shrinkage enabled (apeglm^70^) for the individual pairwise comparisons. An adjusted threshold p-value of 0.05 was applied to the results for generation of upset plots^71^ and Gene expression/LogFC plots. For the generation of ternary plots and subsequent volcano plots further cutoff of absolute log2FC > 2 vs untreated sample group was applied. Gene set enrichment analysis (GSEA)^72^ was conducted on the unfiltered DE comparisons, ranked by shrunken Log2Fold change and using the Hallmark genesets as reference^73^. An adjusted p-value cutoff of 0.05 was applied to the resultant enrichments.

#### Functional enrichment of genes from selected modules and DE genes

Metascape was used to identify biological pathways and processes enriched in the dataset^28^. Genes from the selected 5 positively associated modules and 5 negatively associated modules were used as input to identify enriched terms. Adjusted p-value cut-off of 0.05 was used to filter and sort the enriched terms. Selected enriched terms from the filtered list (that pass adjusted p-value 0.05) from the KEGG pathways and gene ontology biological processes (GO BP) were visualized using barplot. Module eigengene is defined as the first principal component of the expression matrix of the corresponding module.

#### Visualization/comparative analysis using single cell datasets

Genes determined to be upregulated compared to their respective control groups were mapped to an integrated dataset of single-cell RNA data from diverse inflammatory diseases across several organs^5^. A Seurat object^74^ was created from the integrated gene-cell matrix obtained from the authors^5^. Genes found to be elevated in the treated (IFN-γ, IL-32, LPS,) samples compared to their appropriate control samples were included in the genes to be mapped to the single-cell sequencing dataset if their expression could be detected above background in the single-cell sequencing dataset. Seurat’s AddModuleScore function was used to compute an enrichment value for each gene set for each cell. Heatmaps were generated using ggplot^75^ and the viridis package^76^.

## SUPPLEMENTAL INFORMATION

## Excel table titles

**Table S1.** Experimental details and raw cytokine data for the protein and peptide library tested on 4 experimental conditions on human macrophages. Related to Figure 1.

**Table S2.** Experimental details and raw cytokine data for the protein and peptide library validated on a secondary screen on human macrophages. Related to Figure 1.

**Table S3.** GW siRNA primary and secondary screen data from HEK-Blue™ TLR4 reporter cells obtained in Sanford Burnham Prebys Medical Discovery Institute. Related to Figure 4.

**Table S4.** Raw and analysed cytokine data from siRNA hit validation in macrophages. Related to Figure 4.

**Table S5:** Single-cell RNA datasets from multiple inflammatory human diseases across several organs. Related to Figures 6 and 7.

---

**Figure S1.**
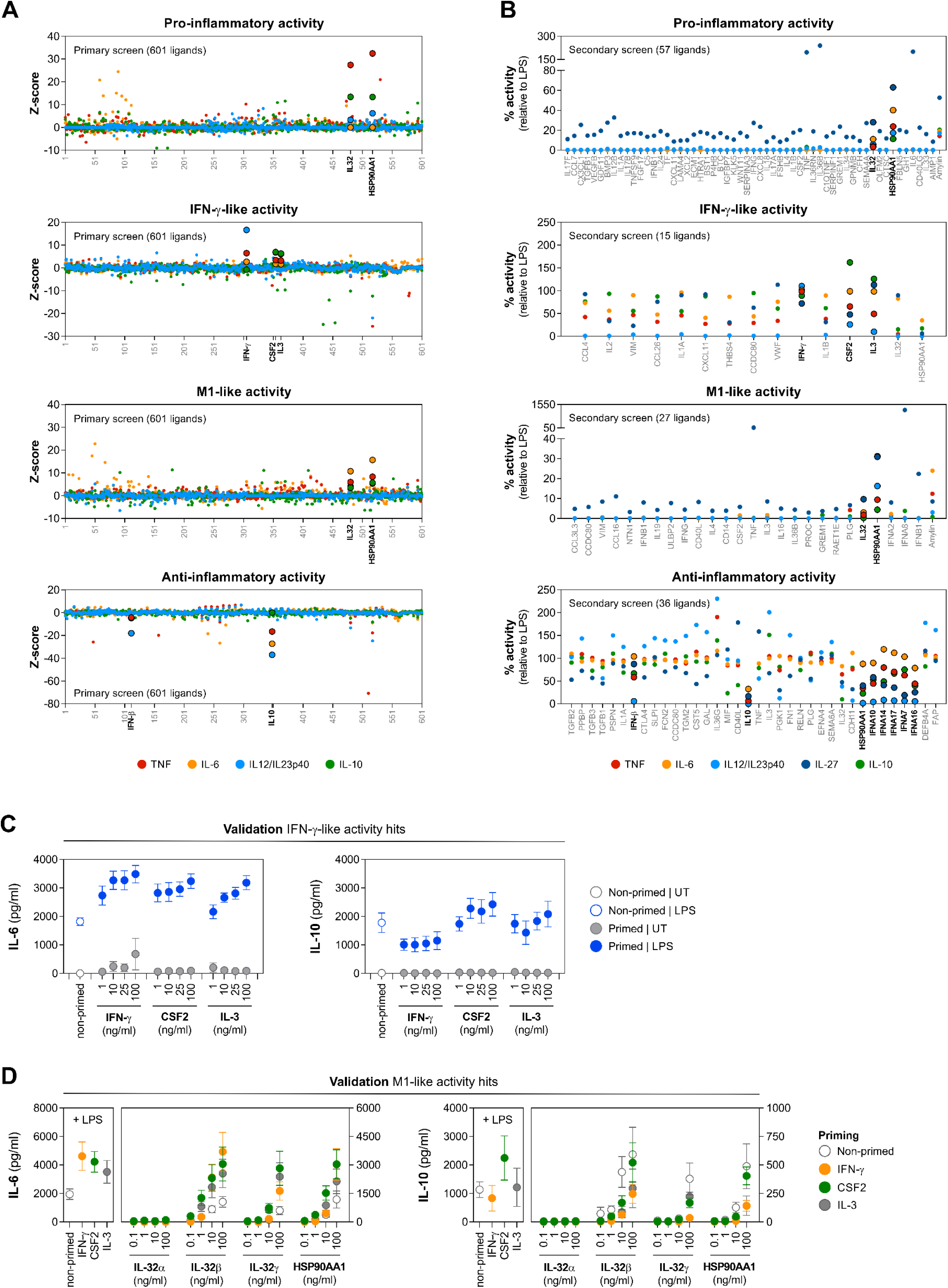
Immunoinflammatory screens in macrophages identify secreted ligands with IFN-γ-like (CSF2, IL-3) and M1-like (IL-32β/γ, HSP90AA1) activities. Related to Figure 1. (A) Global overview of Z-score distribution for all ligands screened in four different phenotypic screens performed in primary human macrophages with a library of 601 recombinant secreted ligands. The most relevant hits which were validated in a secondary screen are highlighted in bold. (B) Validation of selected hits from primary screens and additional selected ligands in a secondary screen, where their activity is represented as relative values to the positive control LPS (50 ng/mL). (C) Validation of macrophage cytokine responses to priming effects of IFN-γ-like ligands (D) Validation of M1-like ligands in non-primed, IFN-γ-primed, CSF2-primed and IL-3 primed macrophages for additional cytokines IL-6 and IL-10.

**Figure S2.**
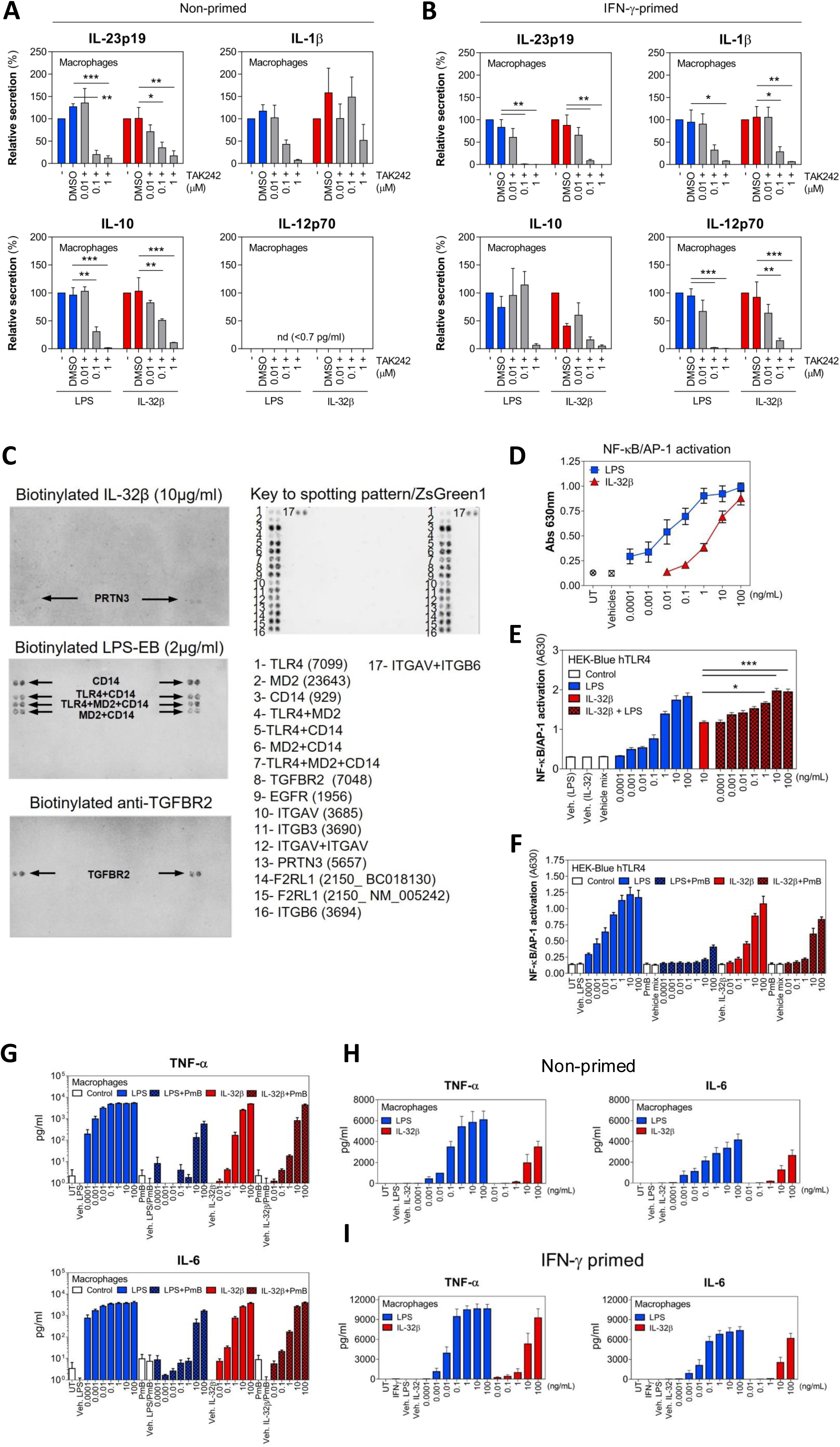
IL-32β inflammatory response is independent of contaminating LPS or direct binding of IL-32β to components of the TLR4 receptor complex. Related to Figure 2. (A-B) Effect of TAK242 on cytokine (IL-23p19, IL-1β, IL-10, IL-12p70) responses from (A) non-primed macrophages or (B) IFN-γ-primed macrophages treated with LPS or IL-32β for 24 h. (C) Binding of biotinylated IL-32β, biotinylated LPS-EB or positive control biotinylated anti-TGFBR2 to receptor expressing HEK293 cells spotted on a cell microarray. HEK293 cells overexpress components of the TLR4 receptor complex (CD14, MD2, TLR4), a positive control (TGFBR2) or candidate receptors for IL-32 from the literature (ITGAV, ITGB3, PRTN3/PAR3, F2RL1, ITGB6). (D) Activation of NF-κB/AP-1 reporter activity by LPS or IL-32β in HEK-Blue™ TLR4 cells treated for 24 h. (E) Concentration-dependent activation of NF-κB/AP-1 reporter by LPS, IL-32β or IL-32β + LPS in HEK-Blue™ TLR4 cells treated for 24 h. (F) Concentration-dependent activation of NF-κB/AP-1 reporter in HEK-Blue™ TLR4 cells treated with LPS or IL-32β +/- the LPS neutralizing antibiotic polymyxin B (PmB, 10 μg/mL) for 24 h. (G) Cytokine (TNF-α, IL-6) responses from non-primed macrophages treated with different concentrations of LPS or IL-32β +/- the LPS neutralizing antibiotic polymyxin B (PmB, 10 μg/mL) for 24 h. (H-I) Cytokine (TNF-α, IL-6) responses from (H) non-primed and (I) IFN-γ-primed macrophages treated with different concentrations of LPS or IL-32β. Statistical analysis: Data shown in (A-B) are mean values ±SEM of n = 3 independent experiments performed on macrophages from one donor, with two-way ANOVA with Dunnett’s multiple comparisons test vs LPS/IL-32β treatment groups (* p < 0.05, ** p < 0.005, *** p < 0.001). Data shown in (E) are mean values ±SEM of 3 independent experiments, with non-parametric Kruskal-Wallis test with Dunn’s multiple comparisons test (*p < 0.05, ***p < 0.001).

**Figure S3.**
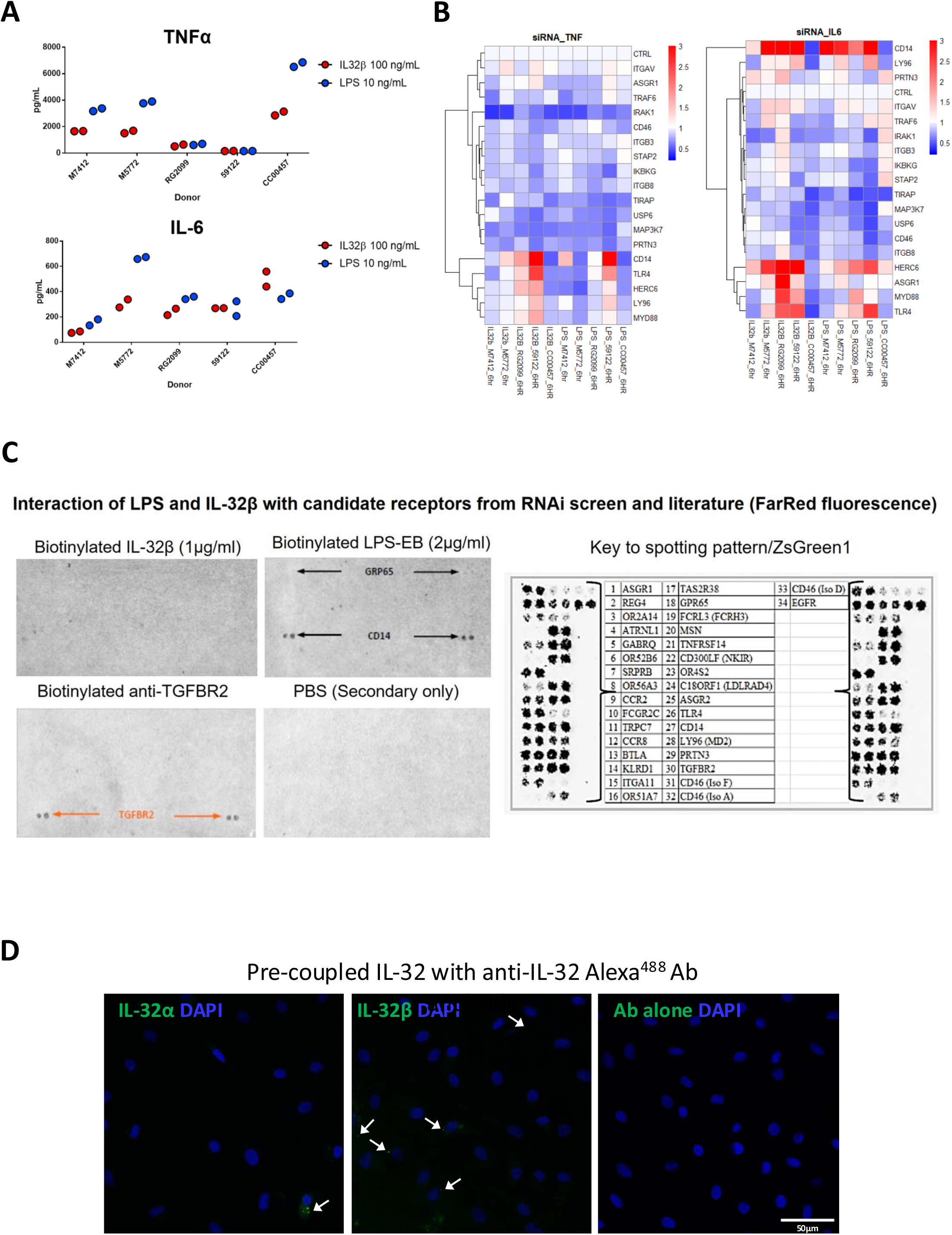
Validation of genome-wide siRNA screen hits identifies IRAK1 as an essential mediator for IL-32β-mediated inflammatory response in macrophages. Related to Figure 4. (A) Donor-specific variation in cytokine (TNF-α, IL-6) responses to LPS (10 ng/mL) or IL-32β (100 ng/mL) from untransfected primary human macrophages used for siRNA experiments (n = 5 donors). (B) Heatmap showing fold change in cytokine (TNF-α, IL-6) responses from macrophages (n = 5 donors) transfected with target-specific siRNA SMARTpools in the validation screen. (C) Binding of biotinylated IL-32β, biotinylated LPS-EB or biotinylated anti-TGFBR2 to HEK293 cells spotted on a cell microarray. Cells overexpress components of the TLR4 receptor complex (CD14, MD2, TLR4), a positive control (TGFBR2) or candidate receptors from the RNAi screen and literature. (D) Representative IF microscopy images of human macrophages incubated for 2 h with pre-coupled IL-32α, or IL-32β (green) to anti-IL-32 Alexa^488^ Ab or Alexa^488^ Ab alone and nuclear staining with DAPI.

**Figure S4.**
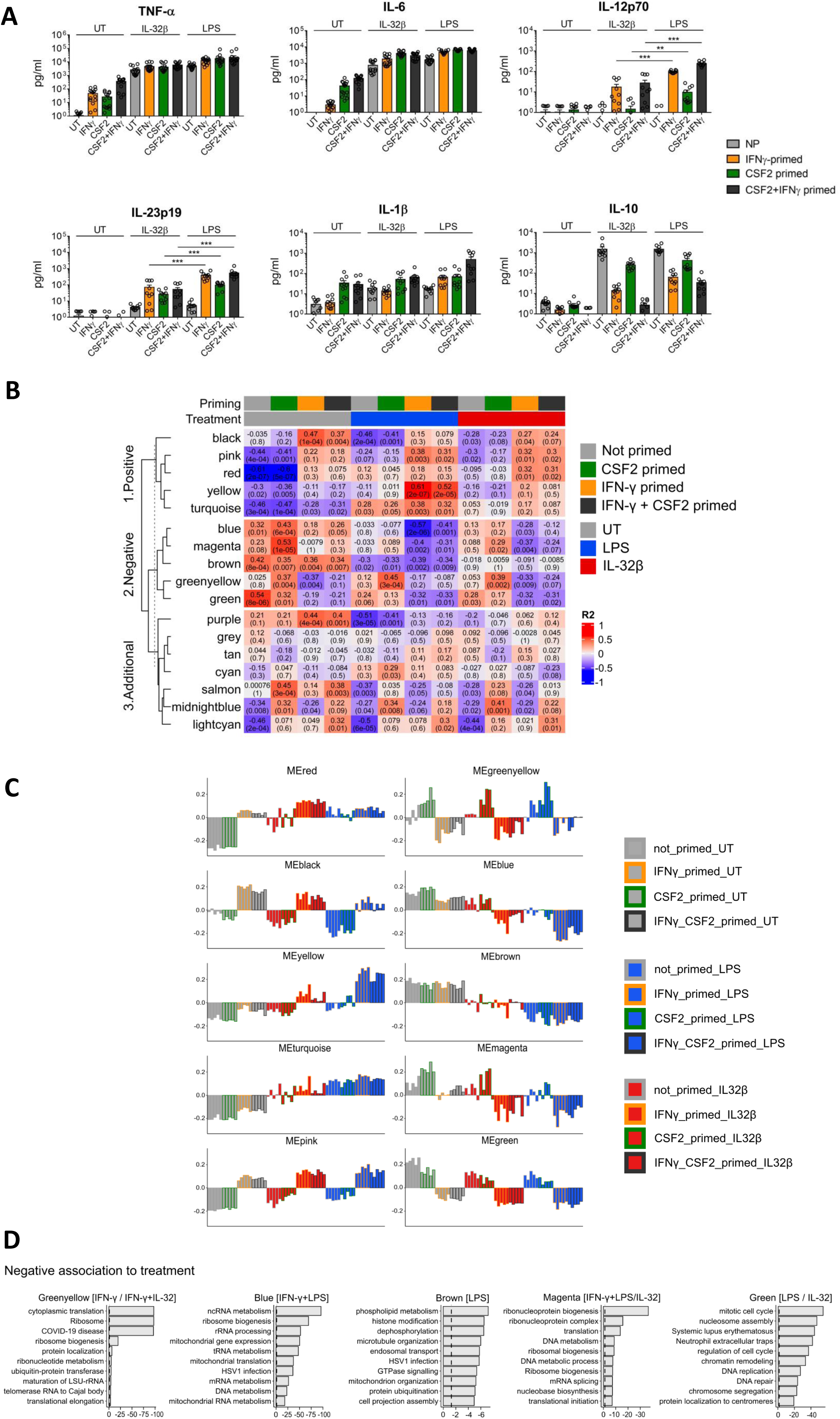
IL-32β induces transcriptional reprogramming of primed and non-primed macrophages with overlapping and distinct gene expression pattern to LPS. Related to Figure 5. (A) Cytokine (TNF-α, IL-6, IL-12p70, IL-23p19, IL-1β, IL-10) responses from primary human monocyte-derived macrophages (n = 5 donors) used for RNA-seq experiments analyzed with LEGENDplex^TM^ assays; untreated and three priming conditions were used for 24 h: IFN-γ primed, CSF2 primed and IFN-γ + CSF2 primed followed by treatment with LPS or IL-32β (10 ng/mL both) for a further 6 h. (B) Module trait relationship heatmap of all modules from WGCNA analyses. The color scale in the heatmap (red to blue) represents the correlation between a module of genes to the trait. Red color indicates positive correlation and blue indicates negative correlation. The top value inside each cell represents R^2^ value representing positive/negative correlation and the bottom value in brackets represents the p-value associated with the significance of the correlation. Modules are assigned a color identifier and row-wise clustered reflecting their positive and negative association to IFN-γ priming treatment and the remaining modules. The top-colored bar indicates the priming status and the colored bar underneath to it indicates treatment groups. (C) Module eigengene bar plot visualization of positive and negative associated modules identified from WGCNA analyses. Module eigengene summarizes the gene expression signatures of entire co-expression modules. It is defined as the first principal component of the expression matrix of the corresponding module. The eigengenes are aligned with along the scaled average expression represented in the y-axis. The fill color in the bar plot represents the treatment and outline color represents the priming group. (D) Bar plots showing the top enriched biological terms (KEGG pathways and GO BP) for gene list in the 5 modules (*greenyellow, blue, brown, magenta, green*) that were negatively associated to IFN-γ priming (IL32/LPS or both) treatment groups as identified by WGCNA analysis. The x-axis in the barplot represents the adjusted p-value of the enriched terms in log scale (-log10(P)). The y-axis represents the selected enriched terms (shortened for aesthetic purposes). Statistical analysis: data shown in (A) are mean values ±SEM of n = 5 independent experiments performed on macrophages from five independent donors, with non-parametric Kruskal-Wallis test with Dunn’s multiple comparisons test (* p < 0.05, ** p < 0.005, *** p < 0.001).

**Figure S5.**
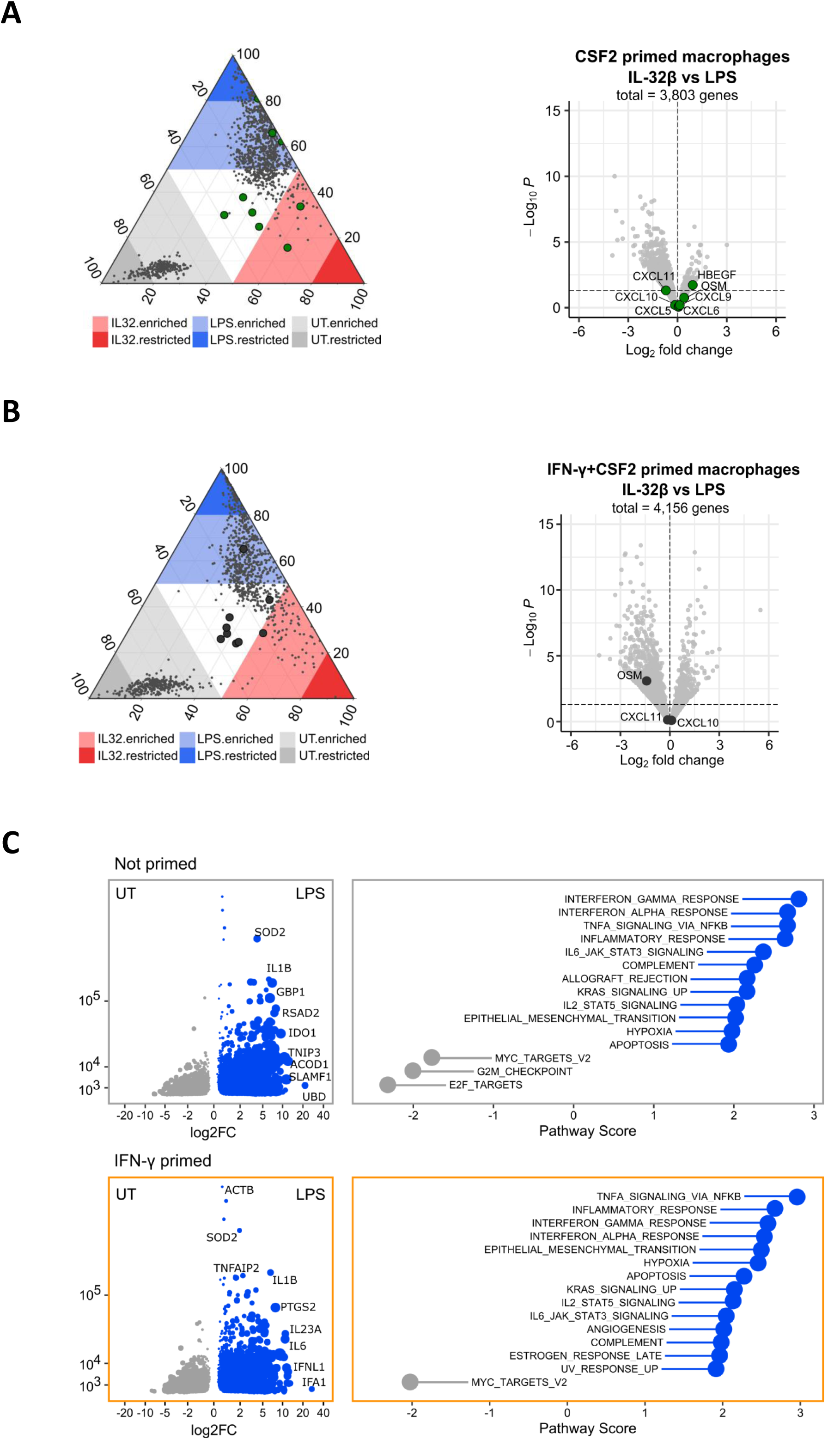
Additional cytokine and transcriptomic data derived from RNA-seq analysis. Related to Figure 5. (A-B) Ternary and volcano plots showing the relative and fold change in gene expression comparing IL-32β and LPS treated human macrophages under (A) CSF2-primed and (B) IFN-γ + CSF2 primed conditions. Genes coding for chemokines selectively induced by IL-32β or LPS in the non-primed condition are shown in both volcano plots. (C) Differentially expressed (DE) genes as a result of LPS treatment in non-primed and IFN-γ primed macrophages are shown with DESea2 base mean parameter as a measure of normalized gene expression (y axis) and shrunken logFC relative to the priming matched UT condition (x axis). Right panel indicates resultant GSEA of the Hallmark gene sets (MSigDB Collections).

**Figure S6.**
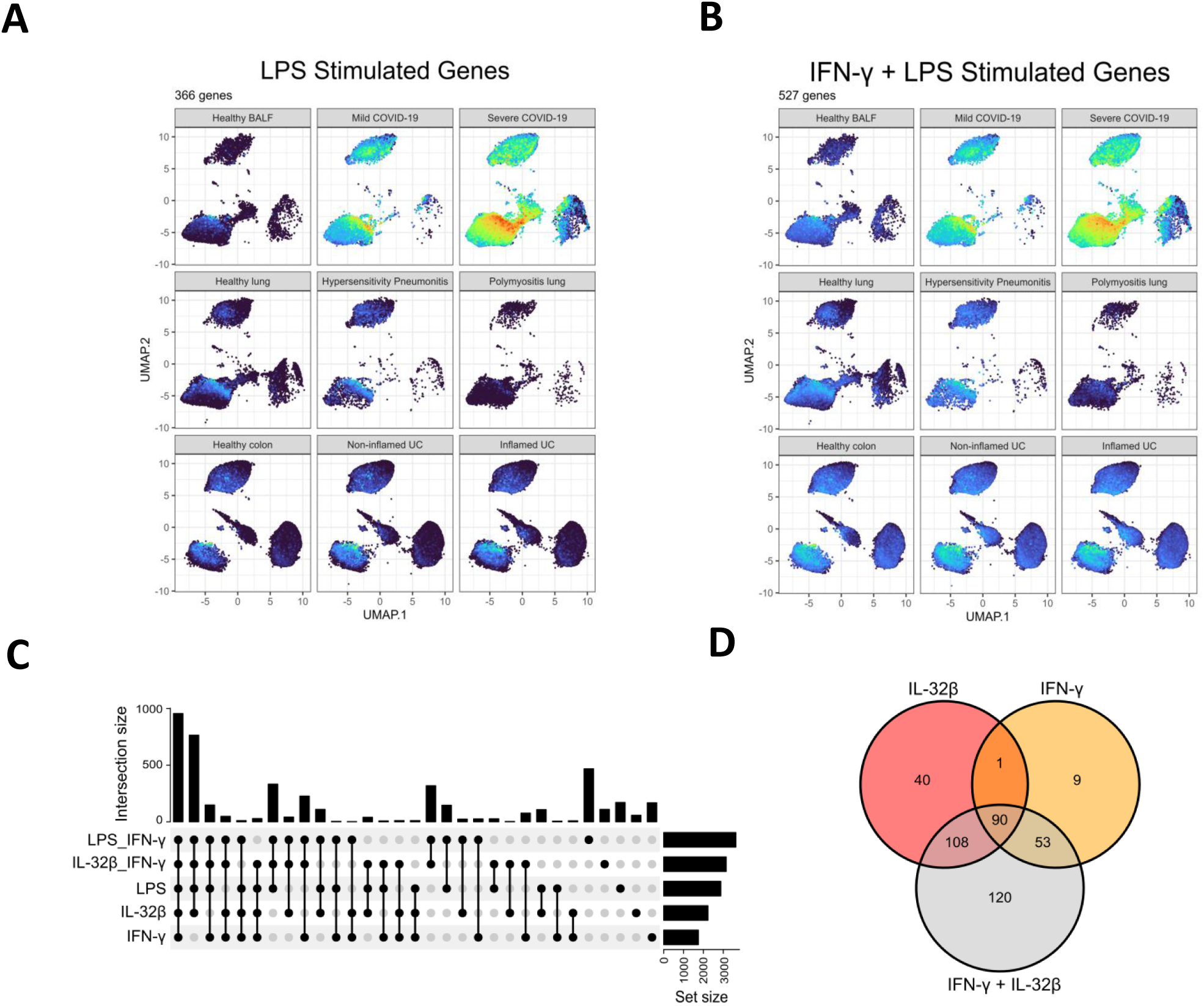
IFN-γ and IL-32β gene expression signatures from human macrophages are enriched in monocyte and macrophage populations in severe COVID-19. Related to Figure 7. (A-B) Expression of LPS and IFN-γ + LPS gene signatures from stimulated human macrophages in single cell populations from multiple inflammatory diseases and COVID-19. Warmer colors indicate a higher degree of overlap. (C) Upset plot showing the number of common and unique genes to each signature. The total number of genes upregulated with each treatment is shown on the right (set size). (D) Venn Diagram showing common upregulated genes to IFN-γ, IL-32β and IFN-γ + IL-32β signatures.

